# A leaky integrate-and-fire computational model based on the connectome of the entire adult *Drosophila* brain reveals insights into sensorimotor processing

**DOI:** 10.1101/2023.05.02.539144

**Authors:** Philip K. Shiu, Gabriella R. Sterne, Nico Spiller, Romain Franconville, Andrea Sandoval, Joie Zhou, Neha Simha, Chan Hyuk Kang, Seongbong Yu, Jinseop S. Kim, Sven Dorkenwald, Arie Matsliah, Philipp Schlegel, Szi-chieh Yu, Claire E. McKellar, Amy Sterling, Marta Costa, Katharina Eichler, Gregory S.X.E. Jefferis, Mala Murthy, Alexander Shakeel Bates, Nils Eckstein, Jan Funke, Salil S. Bidaye, Stefanie Hampel, Andrew M. Seeds, Kristin Scott

## Abstract

The forthcoming assembly of the adult *Drosophila melanogaster* central brain connectome, containing over 125,000 neurons and 50 million synaptic connections, provides a template for examining sensory processing throughout the brain. Here, we create a leaky integrate-and-fire computational model of the entire *Drosophila* brain, based on neural connectivity and neurotransmitter identity, to study circuit properties of feeding and grooming behaviors. We show that activation of sugar-sensing or water-sensing gustatory neurons in the computational model accurately predicts neurons that respond to tastes and are required for feeding initiation. Computational activation of neurons in the feeding region of the *Drosophila* brain predicts those that elicit motor neuron firing, a testable hypothesis that we validate by optogenetic activation and behavioral studies. Moreover, computational activation of different classes of gustatory neurons makes accurate predictions of how multiple taste modalities interact, providing circuit-level insight into aversive and appetitive taste processing. Our computational model predicts that the sugar and water pathways form a partially shared appetitive feeding initiation pathway, which our calcium imaging and behavioral experiments confirm. Additionally, we applied this model to mechanosensory circuits and found that computational activation of mechanosensory neurons predicts activation of a small set of neurons comprising the antennal grooming circuit that do not overlap with gustatory circuits, and accurately describes the circuit response upon activation of different mechanosensory subtypes. Our results demonstrate that modeling brain circuits purely from connectivity and predicted neurotransmitter identity generates experimentally testable hypotheses and can accurately describe complete sensorimotor transformations.

## Main

The *Drosophila* central brain connectome, comprising over 125,000 neurons and 50 million synaptic connections, allows brain-wide analyses of how the fly processes sensory information (Dolan et al., 2014; Ohyama et al., 2015; Huang et al., 2018; Zheng et al., 2018; Meinertzhagen 2018; Bates et al., 2020; Schlegel, P. et al, 2020; Scheffer, L. K. et al., 2020; Marin et al., 2020; Dorkenwald et al., 2022; Lappalainen et al., 2023; Winding et al., 2023; Flywire.ai; forthcoming Flywire paper, 2023). Yet despite the comparative simplicity of the fly brain, a single neuron may be connected to hundreds of downstream neurons, and interpreting connectivity to examine behavior is not straightforward.

To model *Drosophila* sensory processing, we implement a simple leaky integrate-and-fire model using the connection weights derived from the entire *Drosophila* connectome of reconstructed Electron Microscopy (EM) neurons (Gerstner et al., 2014; Kakaria et al., 2017; Pisokas et al., 2020; Churgin et al., 2021; Lazar et al., 2021; Dorkenwald et al., 2022; forthcoming Flywire paper, 2023), as well as neurotransmitter predictions for each neuron (Buhmann et al., 2019; Heinrich et al., 2018; Eckstein et al., 2020). In this model, spiking of a neuron alters the membrane potential of downstream neurons in proportion to the connectivity from the upstream neuron (Gerstner et al., 2014; Lazar et al., 2021; Figure 1A and Methods); if a downstream neuron’s membrane potential reaches the firing threshold, that neuron, in turn, fires. We implemented this model in the spiking neural network simulator Brian2 (Stimberg, Brette, and Goodman, 2019). The baseline firing of each neuron in our model is 0 Hz. By driving activity in a sparse set of neurons, one can predict changes in downstream firing as a result of this activation. Alternatively, one can activate a set of neurons, while also selectively silencing particular neurons, and predict how this silencing alters circuit activity.

**Figure 1.**
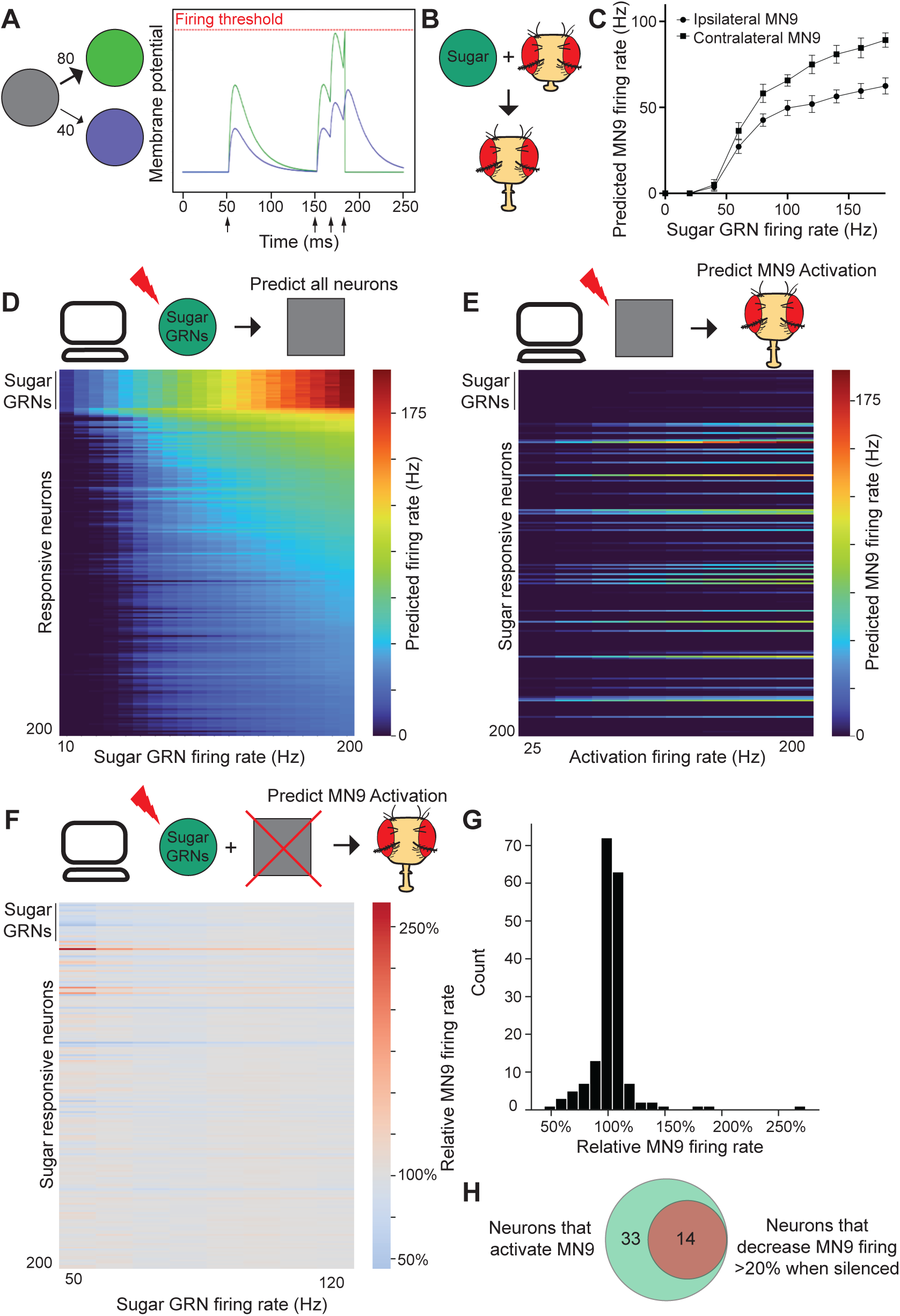
The computational model accurately predicts neurons that respond to sugar stimulation and neurons required for proboscis extension to sugar. A. Schematic of the leaky integrate-and-fire model. Activation of the grey neuron at the times indicated by the arrows results in depolarization of the green and purple neurons in proportion to their connectivity from the grey neuron. When the membrane potential of the green neuron reaches the firing threshold, this neuron fires, and its membrane potential is reset to the resting potential. B. Schematic of the proboscis extension response: presentation of sugar results in exten­ sion of the proboscis. C. Predicted MN9 firing rate of either the ipsilateral or contralateral MN9 in response to unilateral right-hemi­ sphere sugar GRN activation. D. Heatmap depicting the predicted firing rates in response to unilateral 10 to 200 Hz sugar GRN firing. The y-axis is ordered by firing rate at 200 Hz sugar activation, and depicts the top 200 most active neurons. E. Heatmap depicting the predicted MN9 firing rate when the top 200 responsive neurons are activated at 25-200 Hz. F. Heatmap depicting the change in the contralateral MN9 firing rate in response to activation of sugar GRNs at the specified firing rate, while individually silencing each of the top 200 responsive neurons. For E and F, the y-axis is ordered as in D. G. Histogram of the non-GRNs in Fat 50 Hz. H. Venn diagram depicting the intersection between neurons predicted to activate MN9 and neurons predicted to cause a 20% decrease in MN9 firing when silenced.

We examined the ability of this model to predict circuit activity in two systems: feeding initiation and antennal grooming. We began by examining the *Drosophila* feeding initiation circuit because it has well-defined taste sensory inputs and motor outputs that are contained in the central brain connectome, allowing for a complete analysis of the circuit. In addition, extensive experimental analysis has identified a core network driving this behavior, providing a ground truth for computational studies (Dahanukar et al., 2001; Scott et al., 2001; Wang et al., 2004; Dahanukar et al., 2007; Gordon and Scott, 2009; Cameron et al., 2010; Chen et al., 2010; Marella, Mann, and Scott, 2012; Inagaki et al., 2012; Flood et al., 2013; Mann, Gordon, and Scott, 2013; Chu et al., 2014; Inagaki et al., 2015; French et al., 2015; Harris et al., 2015; Miyazaki et al., 2015; Jaeger et al., 2018; Devineni et al., 2019; McKellar et al., 2020; Engert et al., 2022; Shiu, Sterne, et al., 2022). We then assessed the performance of the model in another well-defined circuit, the antennal grooming circuit, as an independent evaluation of the model (Seeds et al., 2014; Hampel et al., 2015; Hampel et al., 2017; Hampel et al., 2020; Zhang, Guo and Simpson, 2020). In both circuits, we tested specific predictions that the computational model generated using cell type specific genetic tools, optogenetics, and functional imaging. We find that the model makes predictions consistent with our empirical observations, such as identification of neurons required for behavioral output. Thus, our computational model reduces the vast complexity of the connectome into simple, intuitive circuits.

In *Drosophila* feeding initiation, detection of appetitive substances in hungry flies results in proboscis extension and consumption (Scott, 2018). Gustatory receptor neurons (GRNs) on the body surface of the fly, including the labellum (the tip of the proboscis) or the legs, directly respond to tastants and project to the primary taste center of the insect brain, the subesophageal zone (SEZ) (Stocker and Schorderet, 1987; Stocker, 1994; Thorne et al., 2004; Miyazaki and Ito, 2010; Hartenstein et al., 2018; Scott, 2018; Engert et al., 2020; Montell, 2021). GRNs respond to specific taste categories, such as appetitive sugar or aversive bitter compounds, resulting in acceptance (i.e., proboscis extension and feeding) or avoidance, respectively (Stocker, 1994; Dahanukar et al., 2001; Dahanukar et al., 2007; French et al., 2015; Montell, 2021; Scott, 2018). We focused on four GRN categories: sugar, water, bitter and putative “low salt” neurons, labeled by Ir94e, to examine the neural circuits that influence feeding in response to taste detection (Jaeger et al., 2018; Scott, 2018; Montell, 2021). These GRNs have been previously identified and classified in the EM brain volume (Engert et al., 2020); we verify and expand on this classification by clustering based on connectivity and comparing this clustering to response properties of second-order neurons (Supplemental Figure 1A-C and Methods).

When a fly encounters sugar, activation of appetitive GRNs results in activation of proboscis motor neurons, leading to proboscis extension and consumption (Figure 1B). In particular, we focused on motor neuron 9 (MN9), which causes extension of the largest portion of the proboscis, the rostrum, when activated (Gordon and Scott, 2009; McKellar et al., 2020). We find that computational activation of labellar sugar-sensing GRNs results in MN9 activation (Figure 1C and Supplemental Figure 1D). Remarkably, unilateral sugar GRN activation more strongly activates the contralateral MN9 compared to the ipsilateral MN9 when either the right (Figure 1C) or the left (Supplemental Figure 1D) hemisphere GRNs are activated. This would be expected to lead to greater extension on contralateral side of the proboscis with the proboscis curving towards the ipsilateral side, consistent with behavioral experiments showing that unilateral taste detection on the legs promotes proboscis extension that is curved and directed toward the food source (Yetman and Pollack, 1987; Schwarz et al., 2017). Thus, we show that *in silico* sensory activation produces activity of motor neurons consistent with the behavior of the fly taste sensorimotor circuit.

We next examined whether computational activation studies would enable us to identify a comprehensive feeding initiation network that included known feeding initiation neurons. We first examined the neural network activated upon unilateral sugar GRN activation. We note that given the variety of assumptions the model relies upon, absolute firing rate predictions are unlikely to be accurate; therefore, we examined network activity upon sugar GRN activation ranging from 10 to 200 Hz (Figure 1D). We find that increasing sugar GRN firing rate not only increases activity of MN9, but also proboscis motor neuron 6 (MN6), which extends the proboscis labella for consumption (McKellar et al., 2020). Of the 127,400 neurons modeled, we found that 45 are predicted to respond to 10 Hz sugar GRN activation, and 455 to 200 Hz (Supplemental Table 1). Activated neurons are defined as neurons that have greater than 0 Hz firing. Thus, the computational model predicts a large network activated by sugar taste detection that includes known sugar-responsive motor neurons.

Sugar taste detection influences activity in nutritive state circuits, memory centers, and modulates a broad range of behaviors, including feeding, oviposition, and foraging (Dethier, 1977; Yang et al., 2008; Scott, 2018; Corfas et al., 2019; Montell, 2021). To evaluate the subset of predicted sugar-responsive neurons that influence feeding initiation, we performed two additional *in silico* experiments. First, as a strategy to identify neurons that drive feeding initiation, we computationally stimulated each of the top responding neurons in the network to identify those that drive activity in MN9 (Figure 1E). Second, to identify neurons required for feeding initiation to sugars, we computationally activated sugar GRNs, silenced each of the top 200 sugar-responsive neurons one at a time, and measured the change in predicted MN9 firing (Figure 1F-G). Neurons that our model predicts to be required for feeding initiation will have decreased MN9 firing when silenced. We defined neurons predicted to cause a silencing phenotype as any neuron whose silencing causes MN9 firing to be 80% or lower compared to control MN9 firing, at any of the 8 sugar activation frequencies tested (Figure 1F). These analyses identified 47 neurons predicted to be sugar-responsive, and sufficient for feeding initiation. Of these 47, 14 neurons are also predicted to be required for MN9 activity (Figure 1H).

We next evaluated whether the predicted neurons for feeding initiation include neurons experimentally shown to participate in feeding initiation behavior. Previous experimental studies of the feeding initiation circuit characterized 10 neural classes that respond to sugar, and are sufficient for proboscis extension (Shiu, Sterne, et al., 2022). Our computational model correctly predicts that all 10 cell types respond to sugar (Supplemental Table 2). Of these 10 neurons, 8 are correctly predicted to be sufficient to activate MN9 (Supplemental Table 2; Shiu, Sterne et al., 2022). We previously found that 5 of the 10 are required for sugar feeding initiation (Shiu, Sterne et al., 2022 and Supplemental Table 2). 3 of these 5 are predicted by our computational model to cause a greater than 20% decrease in MN9 firing, while one of the others is predicted to cause a statistically significant decrease in MN9 firing, but less than 20%, when silenced. Although the model predictions generally match previous experimental results, there are some deviations. For example, two taste-responsive neurons, Usnea and Phantom, are not predicted to activate MN9 (Figure 1D and Supplemental Table 2), however, optogenetic activation of these neurons causes proboscis extension (Shiu, Sterne et al., 2022). These two cell types are predicted to be inhibitory. We note that because the basal firing rate of all neurons in the model is 0, activation of inhibitory neurons in the model, in the absence of other input, cannot alter the firing of downstream neurons. Alternatively, incorrect neurotransmitter predictions (e.g., the neurons are peptidergic or excitatory) or other assumptions of the model may explain discrepancies. Despite these limitations, overall, this analysis demonstrates that this computational approach correctly identifies known neurons in a sensorimotor circuit and suggests additional neurons likely to contribute to this behavior (Supplemental Table 1).

As an independent assessment of whether the computational model accurately predicts neurons that elicit MN9 activity, the output of our sensorimotor circuit, we compared behavioral phenotypes upon optogenetic activation of single cell types with their corresponding computational activation phenotypes. To do this in a non-biased way, we performed a screen where we optogenetically activated split-GAL4 lines and monitored the activity of MN9. The SEZ split-Gal4 collection labels 138 cell types, or an estimated third of all neurons in the SEZ, a feeding region of the brain (Sterne et al., 2020). From the SEZ split-GAL4 collection, we identified 106 cell types in the Flywire volume. We compared the predicted *in silico* MN9 activation phenotypes of these neurons with the actual optogenetic activation MN9 phenotypes we observed. When we activate each cell type at 50 Hz, 11 are predicted to activate MN9; remarkably, 10 of 11 of these cell types actually do elicit MN9 activity when optogenetically activated (Figure 2A-B, and Supplemental Table 3). Furthermore, of the 95 predicted not to elicit proboscis extension due to 50 Hz activation, just 4 have non-zero MN9 extension. Thus, the computational model can predict the activation phenotypes of a non-biased sample of cell types at greater than 90% accuracy.

**Figure 2.**
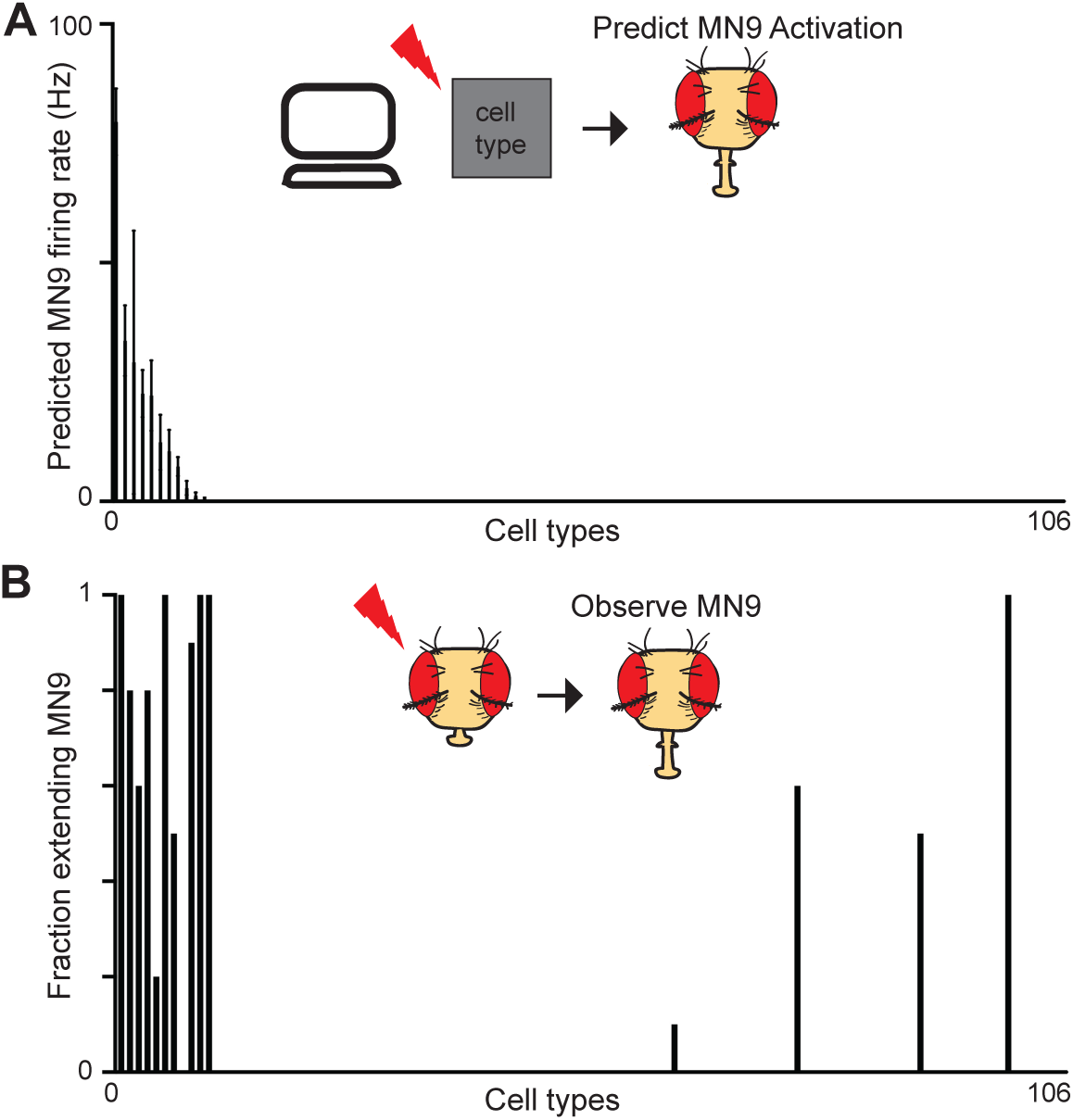
The computational model predicts neurons that elicit MN9 firing. A. Predicted MN9 firing rates when each of 106 cell types are computationally activated at 50 Hz. Cell types are ordered by predicted MN9 firing rate. B. Fraction of flies extending MN9 in response to optogenetic activation. Cell types are ordered as in A. n = 10 flies per cell type.

The accuracy of the model suggests that it provides a powerful platform to query the taste-induced activity of neurons in the brain to discover how different taste modalities are processed to influence feeding initiation. We first tested if the model can predict the response to co-activation of both an attractive sugar stimulus and an aversive bitter stimulus. Bitter detection inhibits proboscis extension motor activity (Meunier et al., 2003). Indeed, the addition of bitter GRN activity to sugar GRN activation in our model resulted in an inhibition of MN6 and MN9 (Figure 3A and Supplemental Table 4). We previously found, using calcium imaging, that bitter GRN activation inhibits the sugar pathway at the level of premotor neurons (Shiu, Sterne et al., 2022), consistent with the predictions of the computational model (Supplemental Table 4).

**Figure 3.**
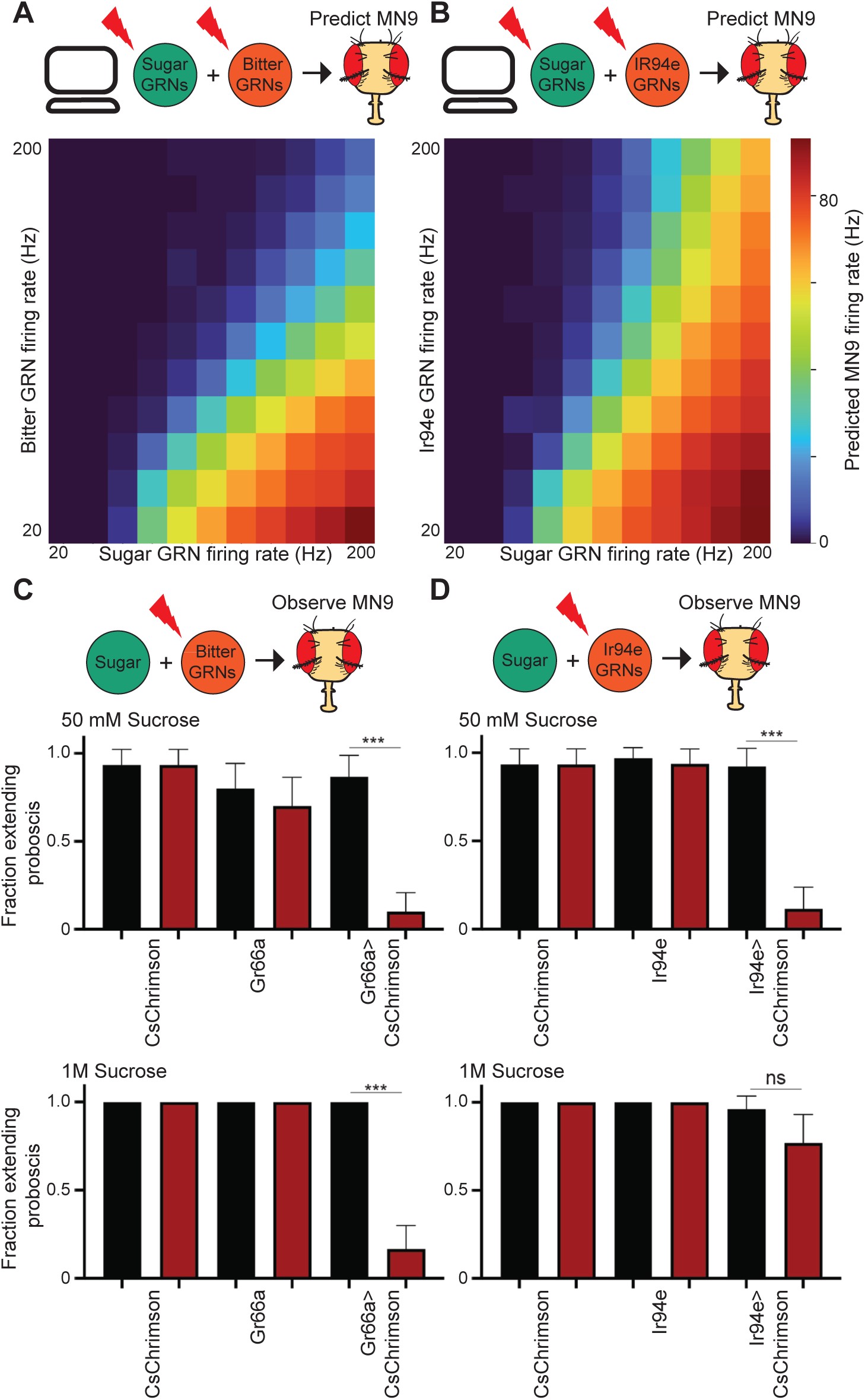
The computational model correctly predicts that lr94e neurons are aversive, but fail to inhibit proboscis exten­ sion to a strong sugar stimulus. A-B. Heatmap depicting the predicted MN9 firing rates in response to the combination of sugar GRN firing and bitter (A) or lr94e (B) GRN activation. C-D. The fraction of flies exhibiting proboscis extension response upon 50 mM sucrose stimulation or 1M sucrose stimulation when Gr66a/bitter GRNs (C) or lr94e/“low salt” GRNs (D) are optogenetically activated. Red bars indicate red light condition. n=26-32. Mean +/- 95% confidence intervals, Fisher’s exact test, ***p<.001.

We next examined the predicted circuit activity caused by GRNs labeled by the ionotropic receptor Ir94e; these neurons have previously been identified in the EM volume (Engert et al., 2020). Ir94e neurons respond to low salt concentrations, among other substances, and are hypothesized to play a role in mediating attraction to low salt (Jaeger et al., 2018). However, the role they play in proboscis extension has not been described. Remarkably, the computational model predicted that activation of Ir94e GRNs, rather than promoting MN9 firing, inhibit MN9 firing (Figure 3B). Therefore, we tested if optogenetic activation of Ir94e GRNs is sufficient to inhibit proboscis extension, similar to bitter activation. Indeed, we found that optogenetic activation of Ir94e GRNs or bitter GRNs was sufficient to inhibit the proboscis extension to 50 mM sucrose, as our modeling predicted (Figure 3C-D). Interestingly, we noted a quantitative difference between the model’s predictions for bitter versus Ir94e activation. Strong bitter activation is predicted to eliminate MN9 firing to strong sugar stimulation, but strong activation of Ir94e neurons is not predicted to do so (see top right portion of 3A-B). We therefore tested the proboscis extension responses to 1M sucrose while optogenetically activating bitter or Ir94e GRNs. Optogenetic bitter activation eliminated consumption of 1M sucrose (Figure 3C), but Ir94e activation did not (Figure 3D). Thus, we conclude that Ir94e GRN activity inhibits proboscis extension, but fails to fully inhibit proboscis extension to strong sugar stimuli, verifying the semi-quantitative predictions of the model and identifying a new aversive taste modality.

Finally, we sought to predict how water taste detection influences feeding initiation. Detection of both sugar and water promote consumption in hungry and thirsty flies, respectively, but the degree to which sugar GRNs and water GRNs activate pathways that are distinct or shared is unknown. We found that activation of water GRNs in our model activates many neurons that are also activated by sugar stimulation. (Figure 4A; Supplemental Tables 1 and 6). In particular, comparing neurons activated by sugar GRNs to those activated by water GRNs, at a stimulation frequency where each pathway activates MN9 at 40 Hz, predicted that the sugar pathway activates 377 neurons, while the water pathway activates 391 neurons. Of these, more than half (250), are shared between the two circuits (Supplemental Figure 2A; Supplemental Table 4). We also examined bitter responsive neurons and Ir94e responsive neurons at the minimal activation sufficient to reduce 40 Hz MN9 firing to 1 Hz. Only 2 neurons were common between sugar and bitter activation, and 30 between sugar and Ir94e activation, demonstrating segregation of neurons activated by aversive and appetitive taste (Supplemental Figure 2A). This prediction is consistent with our previous calcium imaging experiments demonstrating that, across 9 sugar-responsive cell types, 0 respond to a mixture of bitter compounds (Shiu, Sterne et al., 2022). In contrast, our model predicts central neurons that respond to both sugar and water taste activation, as well as sugar-specific and water-specific neurons, consistent with brain-wide calcium imaging studies (Harris, 2015; Münch et al., 2022).

**Figure 4.**
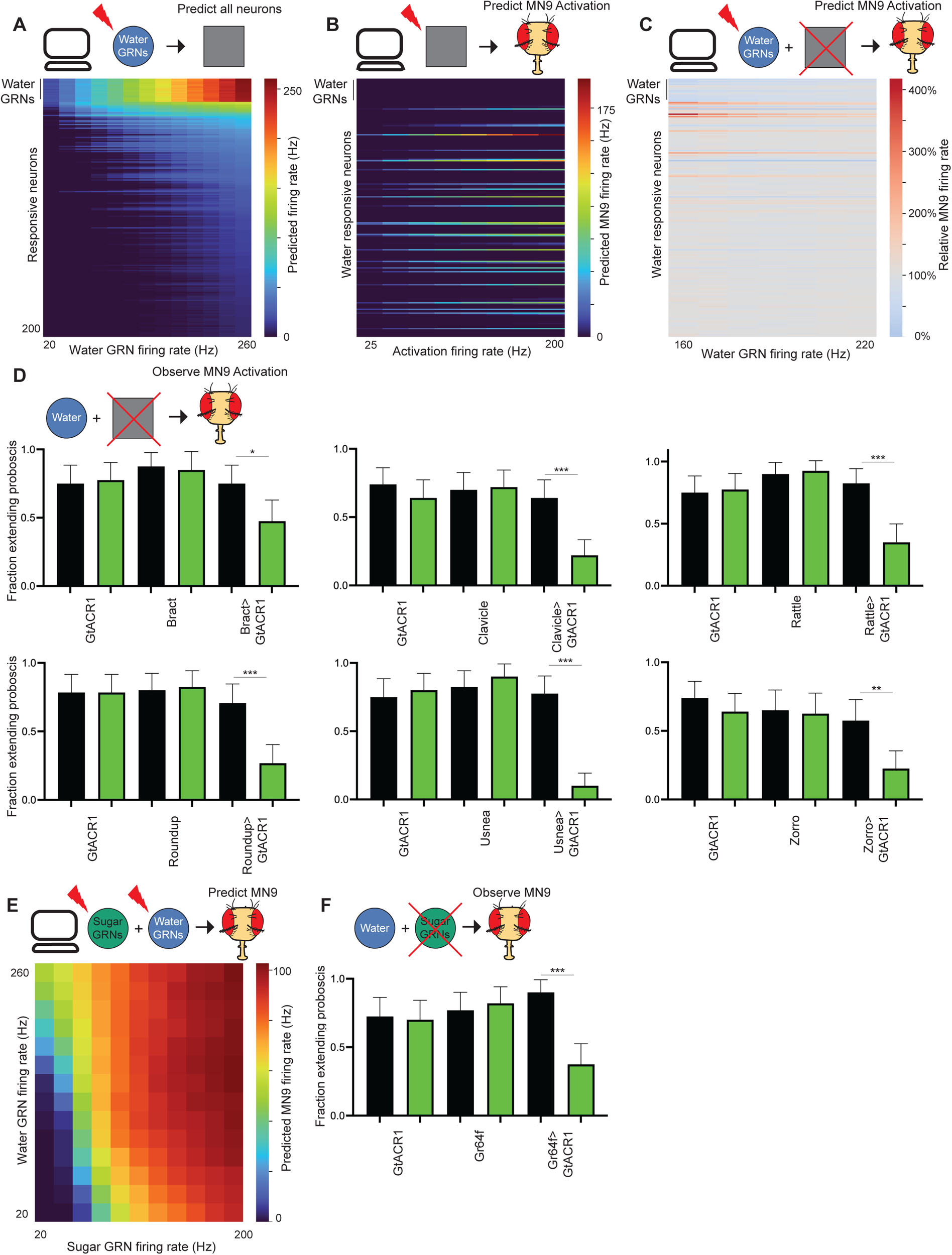
The computational model correctly predicts that the sugar and water pathways share components and additively promote proboscis extension. A. Heatmap depicting the predicting firing rates in response to 20 to 260 Hz water GRN firing. The y-axis is ordered by firing rate at 260 Hz water activation. B. Heatmap depicting the predicted MN9 firing rate when the top 200 respon­ sive neurons are activated at 25-200 Hz. C. Heatmap depicting the change in MN9 firing rate in response to activation of water GRNs at the specified firing rate, while individually silencing each of the top 200 responsive neurons. D. The fraction of flies exhibiting proboscis extension response upon water stimulation. All but Usnea are predicted to cause water silencing phenotypes. Green bars indicate green light condition. n = 30-50. E. Heatmap depicting the predicted MN9 firing rates in response to the combination of sugar and water GRN activity. F. The fraction of flies exhibiting proboscis extension response upon water stimulation. D, F: Mean +/- 95% confidence intervals, Fisher’s exact test, **p<.01, ***p<.001.

To identify interneurons that compose the water feeding initiation circuit, we used the computational model to analyze the water-responsive neurons that influence MN9 activity. We stimulated the top 200 neurons that are predicted to respond to water, and identified the subset that computationally activates MN9, in order to identify water-responsive interneurons that elicit proboscis extension (Figure 4B). Next, we also computationally activated water-sensing GRNs, silenced each water-responsive neuron, and monitored the change in MN9 activity (Figure 4C; Supplemental Figure 2B-C). Using our computational model, we identified 39 water-responsive neurons that are also sufficient for MN9 activation. Of these 39 water-responsive neurons, 30 are also predicted to be activated by sugar GRNs (Supplemental Figure 2B). Furthermore, we identify 9 neurons predicted to be both necessary and sufficient for water feeding initiation (Supplemental Figure 2C). As with sugar, we defined a neuron predicted to be required for water feeding initiation as any neuron that, when silenced, caused MN9 firing to be less than 80% of that of the unsilenced control, at any of the seven frequencies tested.

To experimentally test these predictions, we performed calcium imaging on two neurons predicted to respond to water, Fudog and Zorro. We found that both neurons indeed responded to water (Supplemental Figure 1C and Supplemental Figure 3A). Additionally, we examined six neurons predicted to have water silencing phenotypes. Five of these, when silenced optogenetically, indeed significantly decreased proboscis extension to water, while a sixth, G2N-1, did not (Figure 4D; Supplemental Figure 3B-C). We also examined 5 neurons that respond to computational water activation, but are not predicted to cause a water silencing phenotype. 4 of these 5 neurons did not have a water silencing phenotype, as predicted, although one, Usnea, did significantly decrease proboscis extension when silenced with GtACR1 (Figure 4D).

Our computational model predicts that the water and sugar pathways share a common set of neurons (Supplemental Figure 2A). Do these shared neurons contribute to feeding initiation? Our calcium imaging experiments (Supplemental Figures 1C and 3A) combined with previous experiments (Shiu, Sterne et al., 2022) confirm that 5 neurons predicted to respond to sugar and water do respond to both sugar and water in vivo: Clavicle, Fudog, Phantom, Rattle and Zorro. Moreover, 4 of these neurons had previously been shown to be sufficient for proboscis extension, 3 of which are also required for sugar feeding initiation (Shiu, Sterne et al., 2022). All 3 are among the neurons we found to be experimentally required for feeding initiation to water, as predicted (Figure 4D). Furthermore, the two other cell types we experimentally found to be required for water, Bract and Roundup, are also predicted to respond to both water and sugar (Figure 4D), and have been found to respond to sugar (Shiu, Sterne et al., 2022). However, previous calcium imaging studies did not identify water responses in these two cell types (Shiu, Sterne et al., 2022). This discrepancy may reflect the greater sensitivity of the behavioral silencing experiments compared to calcium imaging experiments of water responses, which are challenging due to the need to bathe the brain in artificial hemolymph, which necessarily alters the thirst state of the fly (Jourjine et al., 2016; Shiu, Sterne et al., 2022). Finally, an additional cell type, Usnea, had been shown to respond to water, but not sugar (Shiu, Sterne et al., 2022); our model correctly predicts Usnea responds to water, but incorrectly predicts that it will also respond to sugar. Usnea has previously been shown to be required for feeding initiation to sugar. We find that it is also required for proboscis extension to water (Figure 4D). Usnea synapses directly onto both sugar and water GRNs (Supplemental Figure 1B), and may tune the response of these neurons. Thus, we identify a set of neurons involved in the processing of both sugar and water.

We therefore next examined the relationship between the water pathway and the sugar pathway by computationally activating both sugar and water GRNs simultaneously and examining the effect on MN9. Our computational modeling predicts that activation of water and sugar GRNs work synergistically to promote MN9 firing (Figure 4F). For example, neither 40 Hz water nor 40 Hz sugar GRN firing is predicted to elicit appreciable MN9 firing, but simultaneous activation of both sugar and water GRNs at 40 Hz is predicted to elicit robust MN9 activity (Figure 4F). If sugar and water do act synergistically, then both sugar GRNs and water GRNs may be involved in water consumption. Only water GRNs have been implicated in proboscis extension to water; we asked if sugar GRNs might also be required. Indeed, optogenetic silencing of sugar GRNs reduced the fraction of flies that elicited proboscis extension to water (Figure 4F). In total, our computational modeling, optogenetic behavior experiments, and functional imaging indicate that the water and sugar pathways share, at least in part, common components to form an appetitive consumption pathway.

To test the general applicability of the computational model to study sensorimotor processing, we sought to determine if it could predict circuit properties in another system. To do this, we focused on the well-studied antennal grooming circuit (Seeds et al., 2014; Hampel et al., 2015; Hampel et al., 2020). In this system, activation of a set of mechanosensory neurons in the Johnston’s organ, a chordotonal organ in the antennae, elicits grooming of the antennae (Kamikouchi et al., 2006, Hampel et al., 2015, Figure 5A). These mechanosensory neurons, abbreviated JONs, synapse onto two interneuron types, named antennal grooming brain interneurons 1 and 2 (aBN1 and aBN2), which in turn synapse onto two descending neurons, aDN1 and aDN2 (Hampel et al., 2015). There is a single aBN1 per hemisphere, while there are multiple aBN2 neurons per hemisphere. Each of these cell types – aBN1, aBN2, aDN1 and aDN2 – are sufficient for antennal grooming, while aBN1 and aBN2 are each at least partially required for antennal grooming (Hampel et al., 2015).

**Figure 5.**
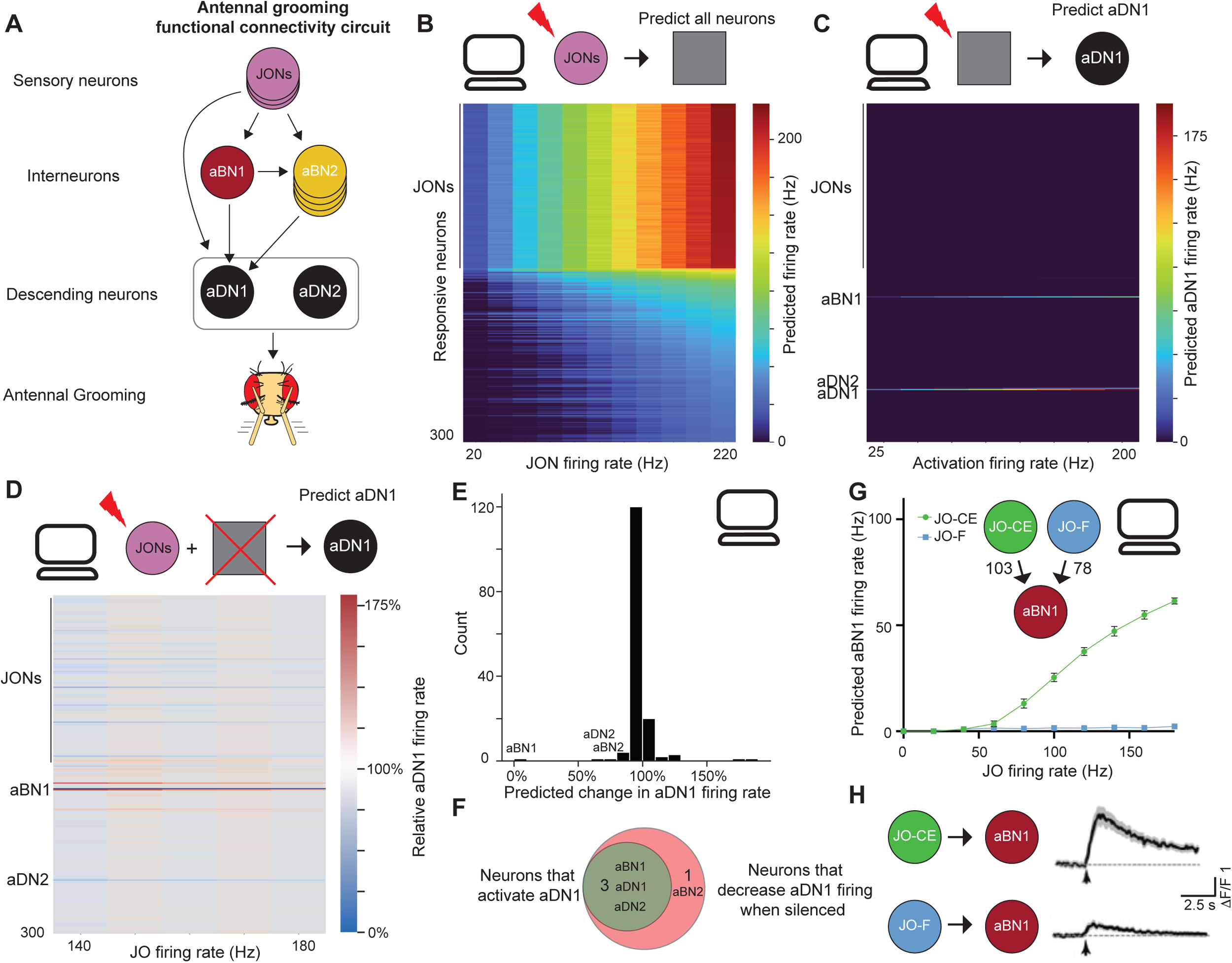
The computational model correctly identifies key neurons in the antenna! grooming circuit as well as subtype circuit responses. A. Schematic of the known antenna! grooming functional connectivity circuit. Arrows represent known functional connectivity (Hampel et al., 2015). Grey oval around aDNs indicates that JONs activate aDNs, but exactly which aDNs are not known. B. Heatmap depicting the predicting firing rates in response to 20 to 220 Hz JON firing. 147 JONs were activated, and are the neurons that have the highest firing rates. The top 300 most responsive neurons are shown. Neurons are ordered by firing rate at 220 Hz C. Heatmap depicting the predicted aDN1 firing rate when the top 300 responsive neurons are activated at 25-200 Hz. D. Heatmap depicting the change in aDN1 firing rate in response to activation of JOs at the specified firing rate, while individually silencing each of the top responsive neurons. E. Histogram of the predicted change in aDN1 firing rate as a result of silencing each non-JONs, when JONs are activated at 140 Hz. Y-axis depicts the number of neurons in each bin. Neurons previously identified are labeled. F. Venn diagram depicting the overlap between neurons predicted to be sufficient to activate aDN1 and neurons required for aDN1 activation. G. JO subtype connectivity onto aBN1 and predicted aBN1 firing in response to JO activation at the specified rate. H. Calcium imaging of aBN1 in response to optogenetic activation of each subtype.

We first sought to test if the computational model could identify the previously described neurons in the circuit. We activated a set of 147 previously identified JONs of the JO-C, JO-E, JO-F, and JO-m subclasses (Hampel et al., 2020). Indeed, the model identified that aBN1, aBN2, aDN1 and aDN2 respond to JON activation (Figure 5B and Supplemental Table 5). To determine which of these JON-responsive neurons might drive antennal grooming, we computationally activated these neurons and asked if they could elicit activity in either of the two descending neurons that evoke antennal grooming, aDN1 or aDN2 (Figure 5C and Supplemental Figure 4A). Next, we asked, among the top neurons predicted to respond to JON activation, which are required for activation of aDN1 or aDN2 (Figure 5D and Supplemental Figure 4B). Remarkably, only four neurons, beyond aDN1 itself, were identified that could elicit aDN1 activity: aBN1, aDN2, and two other neurons that elicited less than 2 Hz aDN1 activity (Figure 5C and Supplemental Table 7). Moreover, only three neurons, besides aDN1 itself, were identified that reduced aDN1 activity by more than 20% at 140 Hz JON activation: aBN1; a descending member of the BN2 class; and aDN2 (Figure 5D-F). Thus, the computational model identifies members of each of the previously identified critical nodes of the antennal grooming circuit, and no additional neurons, purely from knowledge of the sensory inputs and descending outputs.

We next tested how different JON subpopulations influence antennal grooming. JONs send their projections to the antennal mechanosensory and motor center (AMMC) in the ventral brain. JO-C and JO-E neurons respond to antennal vibrations and project medially into the AMMC, while JO-F neurons project into a distinct region (Hampel et al., 2020). Optogenetic activation of both JO-CE and JO-F neurons is sufficient to trigger antennal grooming, but it is not known if these two populations generate distinct patterns of downstream firing. Both JO-CE and JO-F neurons synapse onto aBN1 (103 and 78 synapses, respectively; Figure 5G), raising the possibility that they elicit grooming by activating aBN1.

Interestingly, our computational model predicts that while JO-CE neurons will elicit robust aBN1 activity, JO-F neurons will not, despite synapsing directly onto aBN1 (Figure 5G). To test this prediction, we optogenetically activated each population of JONs and performed calcium imaging in aBN1. Consistent with the model’s prediction, JO-CE robustly activated aBN1, but JO-F neurons did not (Figure 5H). Why do JO-F neurons fail to robustly activate aBN1? We identified three putative inhibitory neurons that are directly postsynaptic to JO-F neurons and synapse directly onto aBN1. Computational silencing of these three neurons permits JO-F neurons to activate aBN1, but this remains to be tested empirically (Supplemental Figure 4C-D).

Our analysis of the antennal grooming circuit demonstrates that our computational model can provide insights into complex circuits, purely from knowledge of sensory input and descending output. Although a naive perspective might suggest that JO-CE and JO-F neurons might activate aBN1 roughly equally because they have approximately similar connectivity onto aBN1, our model accurately predicts aBN1 will robustly respond to JO-CE, but not JO-F activation. In total, we demonstrate that modeling brain circuits purely from connectivity and neurotransmitter identity is sufficient to reliably describe, at least at a coarse level, entire sensorimotor transformations.

In conclusion, we report a computational model, based on the connectivity and neurotransmitter predictions of the entire fly connectome that can predict circuit neural activity, the neurons required for activation of output neurons, and the integration of multiple sensory modalities. We use the model to create predictions of the complete sugar, water, bitter and Ir94e pathways and validate many of these predictions experimentally. We show that the Ir94e neurons, previously hypothesized to be attractive, instead inhibit proboscis extension. Our modeling suggests that sugar, bitter and Ir94e GRNs activate generally distinct populations of neurons. In contrast, sugar and water GRNs activate many of the same central neurons as well as sugar-specific and water-specific neurons. These studies suggest segregated processing of aversive and appetitive tastes, as well as shared and segregated processing for appetitive tastes. In addition, we recapitulate the antennal grooming circuit purely from sensory input and descending output, and identify a subpopulation of JONs that, despite strong connectivity onto aBN1, fail to activate it. These studies demonstrate the power of computational modeling to elucidate sensory processing features in complex networks.

Our analysis of the taste and antennal grooming circuits shows we can model local sensorimotor transformations in the taste and antennal grooming circuits. The computational model, implemented in the widely used Brian2 library (Stimberg, Brette, and Goodman, 2019), allows for perturbations that are easily interpreted, helping to reduce the vast complexity of the connectome into intuitive testable predictions. We believe our computational model will be a useful tool for the study of sensorimotor transformations and the exploration of interactions between overlapping neural pathways (e.g. sweet-bitter, sweet-water, etc). One exciting possibility is modeling interactions between circuits that have generally previously been studied in isolation.

A strength of computational modeling in general is that it is explicit about its assumptions and limitations. In this simple leaky integrate-and-fire model, we treat each neuron identically as a spiking neuron, ignore neural morphology, as well as different neurotransmitter receptor dynamics (Gerstner et al. 2014). Furthermore, the model does not account for the possibility of gap junctions, non-spiking neurons, internal state, or long-range neuropeptides, and assumes that the basal firing of each neuron is 0 (Dermietzel, and Spray, 1993; Rozental, Giaume, and Spray, 2000; Bargmann, 2012, Marder, 2012; Nusbaum, 2017; Nässel and Zandawala, 2019; Flavell, 2022). In addition, the model’s accuracy is limited by the underlying synapse and neurotransmitter prediction accuracy (Buhmann et al., 2019; Heinrich et al., 2018; Eckstein et al., 2020). Moreover, studies of the connectomes of *C. elegans* and the crustacean stomatogastric ganglion demonstrate that connectivity knowledge constrains, but does not dictate, a particular circuit mechanism (Marder, and Bucher, 2007; Bargmann, and Marder, 2013; Scheffer and Meinertzhagen 2021). Despite these limitations, the model performs remarkably well for the demonstrated use cases. Across 164 predictions we were able to test empirically, 91% of predictions were consistent with our empirical results (Supplemental Table 9). Excluding our optogenetic split-GAL4 experiments (Figure 2), in which the vast majority of cell types did not elicit MN9 activation, the model’s accuracy is 84% (Supplemental Table 9). Further refinements of our computational model, for example, more complete neurotransmitter or receptor information, or more sophisticated treatment of the morphology of each neuron, may improve the accuracy of future models. We show here that, in the intermediate complexity of the entire *Drosophila* brain, a simple computational model can reliably describe entire sensorimotor transformations from sensory input to descending or motor output.

## Methods

### Computational Model

We implement a leaky integrate-and-fire model as previously described (Lazar et al., 2021; Kakaria and de Bivort, 2017; Churgin et al., 2021) with α-synapse dynamics, using the following two differential equations and parameters:

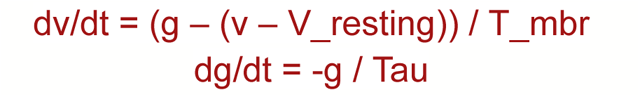

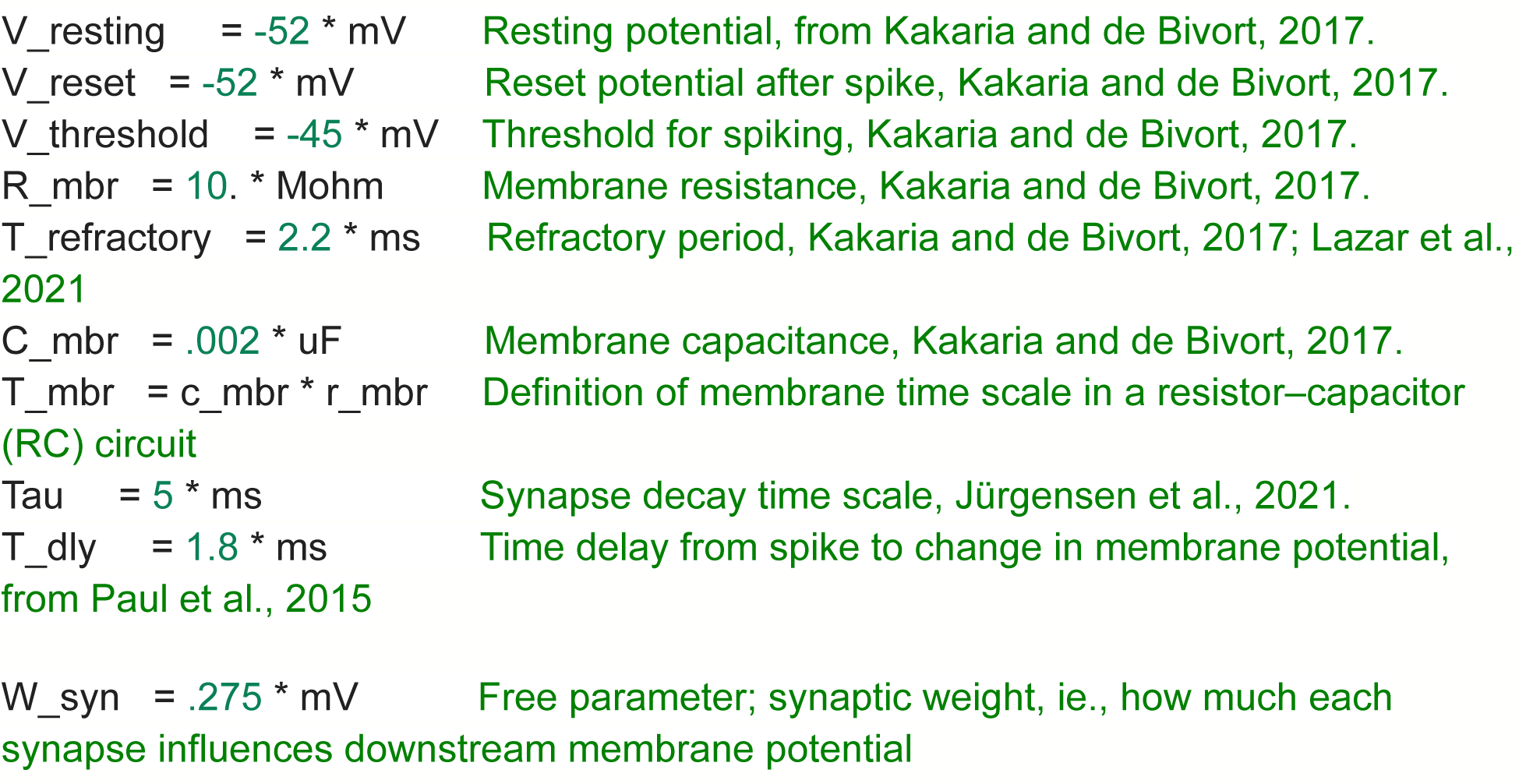

|  |  |
| --- | --- |
| $V_{\text{resting}} = -52 * \text{mV}$ | Resting potential, from Kakaria and de Bivort, 2017. |
| $V_{\text{reset}} = -52 * \text{mV}$ | Reset potential after spike, Kakaria and de Bivort, 2017. |
| $V_{\text{threshold}} = -45 * \text{mV}$ | Threshold for spiking, Kakaria and de Bivort, 2017. |
| $R_{\text{mbr}} = 10. * \text{Mohm}$ | Membrane resistance, Kakaria and de Bivort, 2017. |
| $T_{\text{refractory}} = 2.2 * \text{ms}$ | Refractory period, Kakaria and de Bivort, 2017; Lazar et al., 2021 |
| $C_{\text{mbr}} = .002 * \text{uF}$ | Membrane capacitance, Kakaria and de Bivort, 2017. |
| $T_{\text{mbr}} = c_{\text{mbr}} * r_{\text{mbr}}$ | Definition of membrane time scale in a resistor–capacitor (RC) circuit |
| $\text{Tau} = 5 * \text{ms}$ | Synapse decay time scale, Jürgensen et al., 2021. |
| $T_{\text{dly}} = 1.8 * \text{ms}$ | Time delay from spike to change in membrane potential, from Paul et al., 2015 |
| $W_{\text{syn}} = .275 * \text{mV}$ | Free parameter; synaptic weight, ie., how much each synapse influences downstream membrane potential |

In this model, v, the membrane potential of the neuron, decays back to V_resting, the resting potential, absent any stimulus. If an upstream neuron fires, the membrane potential changes in proportion to the connectivity. If the upstream neuron is excitatory, the neuron depolarizes; if inhibitory, the neuron hyperpolarizes.

All parameters are taken from previous *Drosophila* modeling or electrophysiology efforts (Paul et al. 2015; Lazar et al., 2021; Kakaria and de Bivort, 2017; Churgin et al., 2021; Jürgensen et al., 2021), from the synaptic weights from the Flywire connectome (Dorkenwald et al., 2022; Buhmann et al., 2021;), or from the neurotransmitter predictions (Eckstein et al., 2020), except for W_syn, the single free parameter of the model, which corresponds to how much the downstream membrane potential changes as a result of a single excitatory or inhibitory synapse. We chose W_syn such that activation of sugar GRNs at 100 Hz resulted in roughly 80% of maximal MN9 firing (Dahanukar, et al., 2007 and Inagaki et al., 2012).

The connection weight, w, of any set of two neurons is the synaptic connectivity weight from the Flywire connectivity multiplied by either 1, if the upstream neuron is excitatory or -1, if the upstream neuron is inhibitory, multiplied by W_syn. We used α-synapse modeling as performed previously (Lazar et al., 2021; Stimberg, et al., 2019; https://brian2.readthedocs.io/en/stable/user/converting_from_integrated_form.html), upon firing of the upstream neuron, the neuron specific value, g, becomes g + w. g, upon initialization of the network, or firing of the neuron, starts at 0 mV. Because the membrane potential dynamics are defined by:

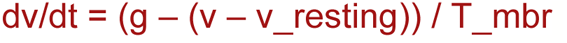

a change in g changes the potential that the neuron will now decay toward.

Furthermore, g exponentially decays with the timescale of tau:

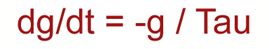

Therefore, after an initial change in membrane potential, once g decays back towards 0, the membrane potential again decays back to the resting potential. Upon firing, a neuron’s membrane potential is reset to the resting potential, and cannot change for the duration of the refractory time period.

We stimulated particular neurons with Poisson distributed input. The neural model is implemented in Brian2 (Stimberg, Brette, and Goodman, 2019), and 30 simulations of 1000 ms for each experiment were performed. All 127,400 proofread neurons from Flywire materialization version 630 are included in the model. The model will be released to a public Github repository upon release of the forthcoming Flywire connectome preprint.

### Neurotransmitter predictions

Neurotransmitter predictions are from Eckstein et al., 2020. In this dataset, the neurotransmitter predictions are predicted for each synapse. We assume GABAergic and glutamatergic neurons are inhibitory (Liu et al., 2013), and that each neuron is either exclusively inhibitory or excitatory. As in Baker et al., 2022, we used a cleft score cutoff of 50, and identified the highest neurotransmitter prediction for each presynaptic site, and if greater than half of all the presynaptic sites across the entire neuron, are predicted to be inhibitory (GABA or Glut), we assigned this neuron as inhibitory.

### Computational modeling limitations

We model neurons as either excitatory or inhibitory spiking neurons, which generally models GABAergic and cholinergic neurons reasonably well, but other neurons (e.g., dopaminergic or serotonergic) will be modeled less well. We assume each neuron is either exclusively inhibitory or excitatory. We ignore neural morphology and receptor dynamics. The underlying synapses or neurotransmitter predictions may not be fully accurate. Gap junctions cannot be identified in the EM dataset, so we ignore their possibility. We do not account for neuropeptides or neuromodulation. Furthermore, we assume a basal firing rate of 0 Hz. An important consequence of this assumption is that inhibitory connections to an inactive neuron have no effect. Because of these assumptions, the absolute firing rates of the model are unlikely to be accurate. Rather, we prefer to interpret changes in firing (e.g., increasing bitter stimulation results in decreasing MN9 firing).

### Computational modeling of water and sugar GRN activity

GRNs were activated at particular frequencies using Poisson distributed input to generate 10-200 Hz firing (sugar) or 20-260 Hz firing (water). Because GRN reconstructions on the right hemisphere are more complete (Engert et al., 2022;), likely due to challenges imaging and reconstructing neurons at the brain’s periphery, we performed unilateral right hemisphere activation for all simulations, except for Supplemental Figure 1D. The firing times of all neurons that fired in any of 30 simulations was recorded, then this data was converted into average firing per second. The top 200 firing neurons were then activated at 25-200 Hz, and the firing of all neurons was recorded. We plot the left hemisphere MN9 firing rate. For computational silencing, water or sugar GRNs were activated, and each of the top 200 firing neurons was silenced by eliminating all output of that neuron. The change in left hemisphere MN9 was plotted.

### Classification of GRNs

Bitter and Ir94e GRNs were previously identified by their distinctive morphology (Engert et al., 2022). Water and sugar GRNs were previously identified by morphology; to confirm these assignments and to incorporate new GRNs identified in Flywire, we performed hierarchical clustering using the connectivity of GRNs onto second-order neurons, using cosine distance as a measure of similarity (Li et al., 2020). This identified three broad clusters of GRNs (Supplemental Figure 1A). We compared this clustering with the response properties of second-order neurons in starved flies, the nutritional state for which we have the most complete data (Shiu, Sterne et al., 2022). Cluster 2 GRNs synapse onto, among others, G2N-1 and FMIN, which respond exclusively to sugar. Cluster 3 neurons synapse comparatively strongly onto Fudog and Phantom, which respond to both water and sugar in starved flies; cluster 3 neurons also, on average, synapse more strongly onto Usnea than cluster 2 neurons. Usnea responds exclusively to water. Thus, we conclude that cluster 2 corresponds to sugar GRNs, consistent with their previous assignment (Engert et al., 2020), and cluster 3 corresponds to water GRNs. We speculate that cluster 1 GRNs are Ppk23-glut GRNs (Jaeger et al., 2018) that may respond to salt, however, little is known about this pathway, making testing this hypothesis challenging. We also examined the connectivity of “zero-order” gustatory neurons onto cluster 2 and 3 GRNs. This clustering corresponds strongly with the GRN to second order clustering (Supplementary Figure 1B), verifying that these clusters reflect distinct taste modalities. As a conservative measure, we chose to examine only those sugar GRNs that fully correspond between the GRN to second order clustering and the zero-order to GRN clustering.

### Identification of SEZ split-GAL4 neurons in the Flywire volume and computational activation

106 cell types, out of 138, of the Sterne et al. SEZ split-GAL4 library were identified in the Flywire volume. Two semi-independent methods were used to identify these neurons:

In the first method, aligned JRC2018 unisex registered brains were downloaded from https://splitgal4.janelia.org, and, where possible, single neuron multicolor Flp-out (MCFO) imagery was used. Neurons were skeletonized in FIJI by selecting a threshold that eliminated the background, but retained the morphology of the neuron. Neurons were skeletonized using the “Skeletonize 2D/3D” tool, and files were saved as a .nrrd. These .nrrd files were converted to .swc in natverse (Bates et al., 2021), and uploaded to the Flywire gateway (https://flywiregateway.pniapps.org/upload; Dorkenwald et al., 2022). This generated pointclouds which were used to manually identify the neuron of interest.

In the second method, we identified the SEZ interneurons in Flywire with the aid of NBLAST (Costa et al., 2016). The morphology of SEZ split-GAL4 interneurons from MCFO images were provided in the dotprops format (Sterne et al., 2021), the points and tangent vectors representing the arborization, from which the pointcloud of each neuron was extracted. The pointcloud was transformed from the JRC2018U brain to FlyWire brain (FAFB) using the Navis and FlyBrains libraries. To narrow down the candidate neurons in FlyWire near the pointcloud, we assigned 1∼4 3D boxes surrounding the dense regions of the pointcloud. The neuronal segments in the boxes were queried using the CloudVolume input/output interface of FlyWire. The skeletons of neuronal segments were calculated and their similarities to the dotprops of the SEZ interneuron were measured using NBLAST. The candidates with the highest NBLAST scores were visually compared to the pointcloud in the 3D view of FlyWire for final decision.

SEZ split-GAL4 neurons were identified independently between the two methods and sets of researchers, and only those for which there was a clear consensus were used. All identified neurons in a cell type were activated computationally, and the resulting MN9 firing rate was recorded.

### Modeling of the antennal grooming circuit

147 JONs of classes JO-C, JO-E, JO-F and JO-mz were previously identified and described in the electron microscopy volume (Hampel et al., 2020). As in the feeding initiation circuit, JONs were activated to generate 20-220 Hz firing. The firing times of all neurons that fired in any of 30 1000 millisecond simulations was recorded, then this data was converted into average firing per second. The top 300 firing neurons were then activated at 50, 100, 150 and 200 Hz to determine if they could activate aDN1 or aDN2. Each of the 147 JONs were activated, and each of the 300 top firing neurons was silenced by eliminating all outputs of that neuron, and the activity of aDN1 or aDN2 was recorded. Each of the aBN1, aBN2, aDN1 and aDN2 were identified from their previously described morphology (Seeds et al., 2015)

### Chrimson Optogenetic activation experiments

Proboscis extension response was scored as previously described (Shiu, Sterne et al., 2022; Mann et al., 2013). Female flies were raised on standard cornmeal-yeast-molasses medium. 48 hr before experiments, flies were placed on molasses food with 0.4 mM retinal. Three- to five-day-old flies were anesthetized with carbon dioxide, mounted onto a glass slide with nail polish, and allowed to recover for 2 hr in a humidified chamber at 22°C. For optogenetic activation experiments, 153 uW/mm^2^ 635 nm laser light was used (Laserglow). Flies were scored for whether they extended their proboscis within a 5 s period in response to light. Experiments were performed blind to genotype. For the screen, 10 flies per genotype were scored for any movement of the proboscis. Split-GAL4s with any extension were scored a second time, from a second, independent cross, specifically for extension of the rostrum, i.e., MN9 activation. If multiple split-GAL4 lines were tested, the split-GAL4 line with the highest MN9 is displayed in Figure 2B.

### GtACR1 silencing

Three-day-old female flies were raised on standard cornmeal-yeast-molasses media, and transferred to standard food with 0.4 mM all-trans retinal for 48 hours. Flies were anesthetized with carbon dioxide, mounted onto a glass slide with nail polish, and and dessicated for 3 hours in a sealed chamber with ∼250g CaSO4 (Drierite, stock# 23001) at 22°C (Jourjine et al., 2016). A green laser (532 nm, LaserGlow LBS-532) was used to acutely silence neurons using GtACR1 (Mohammad et al., 2017). Flies presented with water three times to the proboscis, and the number of flies that extended at least once was recorded.

### Ir94e and Gr66a optogenetic activation

Experiments were performed as in the GtAcr1 experiments, except flies were exposed to red light, rather than green light. Flies were raised on 0.4 mM retinal in standard food for 4 days. Flies were anesthetized with carbon dioxide, mounted onto a glass slide with nail polish, and allowed to recover for 2 hr in a humidified chamber at 22°C. 153 uW/mm2 635 nm laser light was used (Laserglow). Flies were water satiated, then presented with either 50 mM sucrose or 1M sucrose three times to the proboscis, and the number of flies that extended at least once was recorded. Experiments were performed blind to genotype.

### Calcium imaging setup for Fudog and Zorro imaging

Mated female flies were dissected for calcium imaging studies 14–21 days post-eclosion as previously described (Shiu Sterne et al., 2022; Harris et al., 2015). Flies were briefly anesthetized with ice as they were placed in a custom plastic holder at the cervix to isolate the head from the rest of the body. The head was then immobilized using UV glue, and the esophagus was cut to provide unobstructed imaging access to the SEZ. For Fudog, flies were food-deprived in a vial containing a wet kimwipe for 18–24 hr prior to imaging. Following dissection, samples were bathed in ∼250 mOsmo AHL and imaged immediately. To generate thirsty-like (pseudodessicated) Zorro flies (Shiu, Sterne et al., 2022), following dissection, samples were bathed in ∼350 mOsmo AHL (‘high osmolality artificial hemolymph’) and allowed to rest for 1 hr prior to imaging.

### Calcium imaging of Fudog and Zorro

For imaging responses to taste solutions, females of UAS-CD8-tdTomato;20XUAS-IVS-GCaMP6s(attP5);20XUAS-IVS-GCaMP6s(VK00005) were crossed to males for each split-GAL4 line, and female progeny without balancers were selected for imaging. The following tastants were used: double-distilled water (‘water’), 1 M sucrose (‘sugar’), or 10 mM denatonium plus 100 mM caffeine in 20% polyethylene glycol (PEG) (‘bitter’). Taste solutions were delivered to the proboscis using a glass capillary (1.0 mm OD/ 0.78 mm ID) filled with ∼4 µL of taste solution and positioned at the tip of the proboscis using a micromanipulator. Taste solutions were drawn away from the tip of the capillary at the beginning of each imaging trial using slight suction generated by an attached 1 mL syringe, and delivered to the proboscis at the relevant time during imaging with light pressure applied to the syringe.

1-photon imaging of Fudog was performed as previously described (Shiu, Sterne et al., 2022) using a 3i spinning disc confocal microscope with a piezo drive and a 20 × water immersion objective (NA = 1.0) with a 2.5 × magnification changer. 55 frames of 8 z sections spaced at 1 µM intervals were binned 4 × 4 and acquired at 0.8 Hz using a 488 nm laser. Taste solutions were in contact with the proboscis labellum from frame 20 to frame 25.

2-photon imaging of Zorro was performed as previously described (Shiu, Sterne et al., 2022) using a Scientifica Hyperscope with resonant scanning, a piezo drive, and a 20× water immersion objective (NA = 1.0) with 4× digital zoom. 80 stacks of 20 z sections spaced at 2 µM intervals were acquired at 0.667 Hz using a 920 nm laser. Taste solutions were in contact with the proboscis labellum from frame 30 to frame 40.

### Functional connectivity between JON subpopulations and aBN1

This experiment required the use of two binary expression systems (GAL4/UAS and LexA/LexAop) for driving the expression of CsChrimson and GCaMP6 in different neurons in the same fly. For the experiment shown in Figure 5H, CsChrimson was expressed in the JON subpopulations using LexA and spGAL4 driver lines that were specific for either JO-CE or -F neurons. LexA and spGAL4 driver lines that were used for expressing GCaMP6 in aBN1 (aBN1-spGAL4-1) were identified in our previous study (Hampel et al., 2015). To identify a driver line that expresses in JO-CE neurons, we screened through the image database described above to identify a LexA driver line, 100C03-LexA (Tirián and Dickson, 2017).

### Taste-response calcium imaging analysis

For calcium imaging analysis of Supplemental Figure 1C and Supplemental Figure 3A, image analysis was carried out in Fiji (Schindelin et al., 2012), CircuitCatcher (a customized Python program by Daniel Bushey Dag et al., 2019), Python, and R. First, in Fiji, Z stacks for each time point were maximum intensity projected and then movement corrected using the StackReg plugin with ‘Rigid Body’ or ‘Translation’ transformation (Thévenaz et al., 1998). Next, using CircuitCatcher, an ROI containing the neurites of the cell type of interest was selected along with a background ROI, and average fluorescence intensity for each ROI at each timepoint was retrieved. Then, in Python, background subtraction was carried out for each timepoint (Ft). To calculate Finitial, initial fluorescence intensity was calculated as the mean corrected average fluorescence intensity from frame 9–18 (for 1-photon imaging) or frame 0–19 (for 2-photon imaging and optogenetic imaging). Finally, the following formula was used to calculate ΔF/F: Ft-Finitial/Finitial. Area under the curve was approximated with the trapezoidal rule in Python using the NumPy.trapz function. Area under the curve was assessed from frames 20–25 (for 1-photon imaging).

## Author Contributions

Philip Shiu conceived the project, wrote the computational model, performed most experiments, and co-wrote the manuscript under guidance from Kristin Scott. Gabriella Sterne performed calcium imaging experiments of Fudog and Zorro, and edited the manuscript. Nico Spiller edited and co-wrote the computational model, and edited the manuscript. Romain Franconville performed calcium imaging experiments of the JON/aBN1 functional connectivity. Philip Shiu, Andrea Sandoval, Chan Hyuk Kang, Seongbong Yu and Jinseop Kim contributed to the identification and annotation of the Sterne et al. split-GAL4 SEZ neurons. Joie Zhou and Neha Simha performed optogenetic activation experiments. Sven Dorkenwald contributed to Flywire infrastructure and management. Arie Matsliah generated the Flywire Codex and Gateway. Philipp Schlegel, Szi-chieh Yu, Claire McKellar, Amy Sterling, Marta Costa, and Katharina Eichler contributed to Flywire community training, proofreading and management. Alexander Bates, Philip Schlegel, Katharina Eichler and Gregory Jefferis contributed to Flywire annotations and cell type matching. Gregory Jefferis led the Cambridge Flywire team. Mala Murthy led the Flywire effort. Alexander Bates, Nils Eckstein, Greg Jefferis and Jan Funke created the brain-wide neurotransmitter predictions. Salil Bidaye provided expertise on the computational model and edited the manuscript. Stefanie Hampel and Andrew Seeds led the antennal grooming portion and edited the manuscript. Kristin Scott co-wrote the manuscript and supervised the project.

## Acknowledgements

We thank members of the Scott Lab for their contributions to the experimental and model design, data analysis and manuscript preparation. We thank Christine Liu, Michael Levy and Gautam Agarwal for their feedback on the initial development of the computational model. We thank the laboratories of Carlos Ribeiro and Rachel Wilson for Flywire proofreading contributions. We thank the entire Flywire community for their contributions to the proofreading of the *Drosophila* connectome. This work was supported by NIH R01DC013280 (KS), NIH F32DK117671 (GRS) and NIH F32DC018225 (PS). Nico Spiller is funded by the Carl Angus DeSantis Foundation. We thank Marissa Sorek for assistance with Flywire community management. We thank Ran Lu, Thomas Macrina, Kisuk Lee, J. Alexander Bae, Shang Mu, Barak Nehoran, Eric Mitchell, Sergiy Popovych, Jongpeng Wu, Zhen Jia, Manuel Castro, Nico Kemnitz, Dodam Ih for alignment and segmentation of the FAFB EM volume and registration to the original FAFB EM dataset. We thank Davi Bock and Eric Perlman for sponsorship of partial proofreading and registration service. We thank Forrest Collman, Casey Schneider-Mizell, Chris Jordan, Derrick Brittain, Akilesh Haligeri for CAVE development and maintenance. We thank Kai Kuehner, Oluwaseun Ogedengbe, Jay Gager, Will Silversmith, Ryan Morey for Neuroglancer development, tools, and Codex development. We thank Sebastian Seung for his suggestions and contributions to Flywire. Flywire is supported by NIH BRAIN Initiative grants MH117815 and NS126935 to Murthy and Seung. Additional proofreading and infrastructure was supported by Wellcome awards 203261/Z/16/Z and 220343/Z/20/Z to Jefferis, and NIMH BRAIN Initiative award 1RF1MH120679-01 and NSF NeuroNex award DBI-2014862 to Bock and Jefferis.

---

**Supplemental Figure 1.**
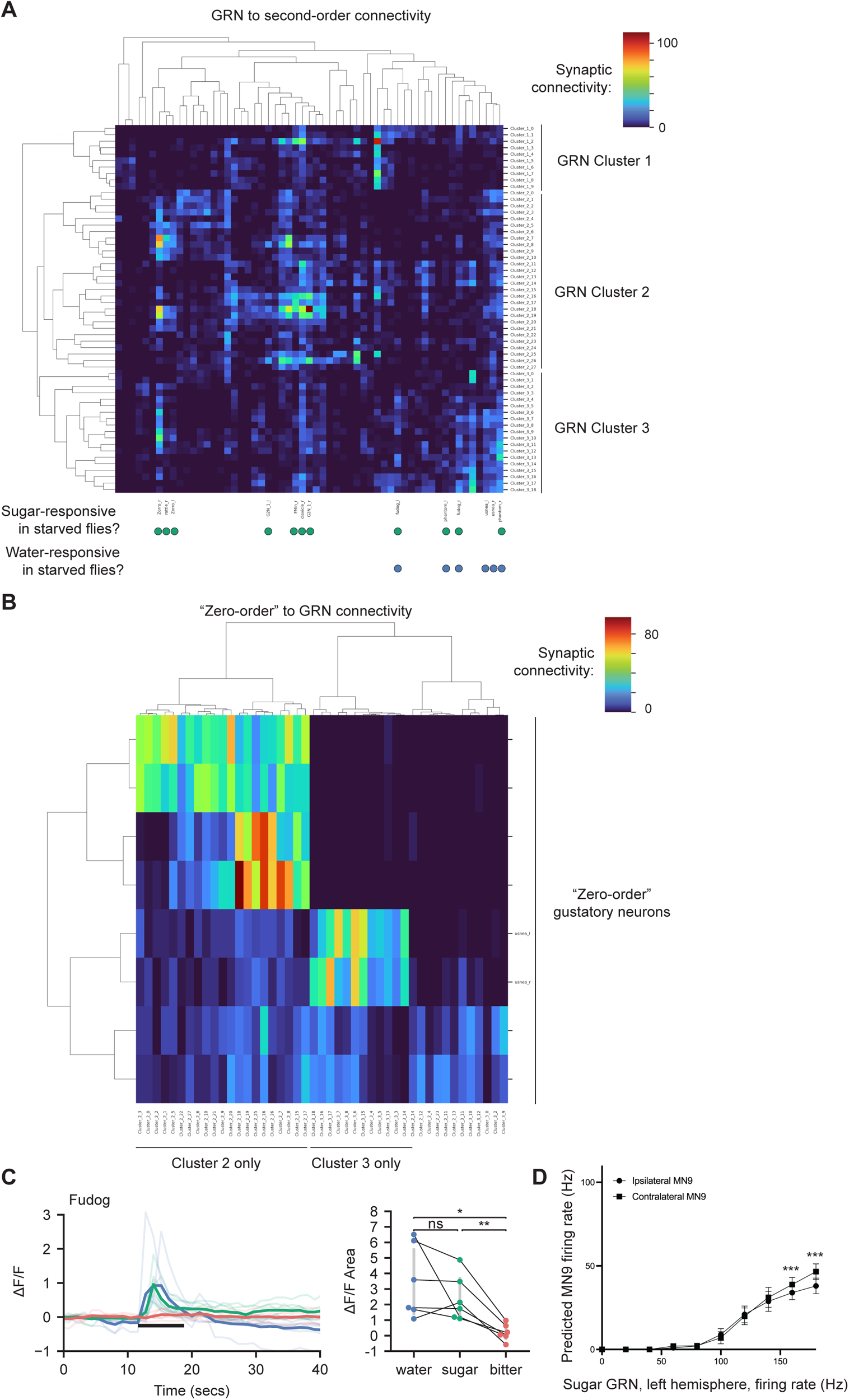
Identification of GRN modality. A. Hierarchical clustering of GRNs and second-order gustatory neurons based on cosine similiarity of connectivity of GRNs (y-axis) onto second-order neurons (x-axis). The response properties of tested second-order neurons in starved flies is plotted (Shiu, Sterne et al., 2022). B. Hierarchical clustering of “zero-order” gustatory neurons onto cluster 2 and 3 GRNs based on cosine similiarity. C. Calcium responses of the second-order neuron Fudog to stimu­ lation of the proboscis in food-deprived flies. D. Predicted MN9 firing rate of either the ipsilateral or contralateral MN9 in response to unilateral left-hemisphere sugar GRN activation. Mean +/- Standard deviation, Mann-Whitney U-test, *** p<.001.

**Supplemental Figure 2.**
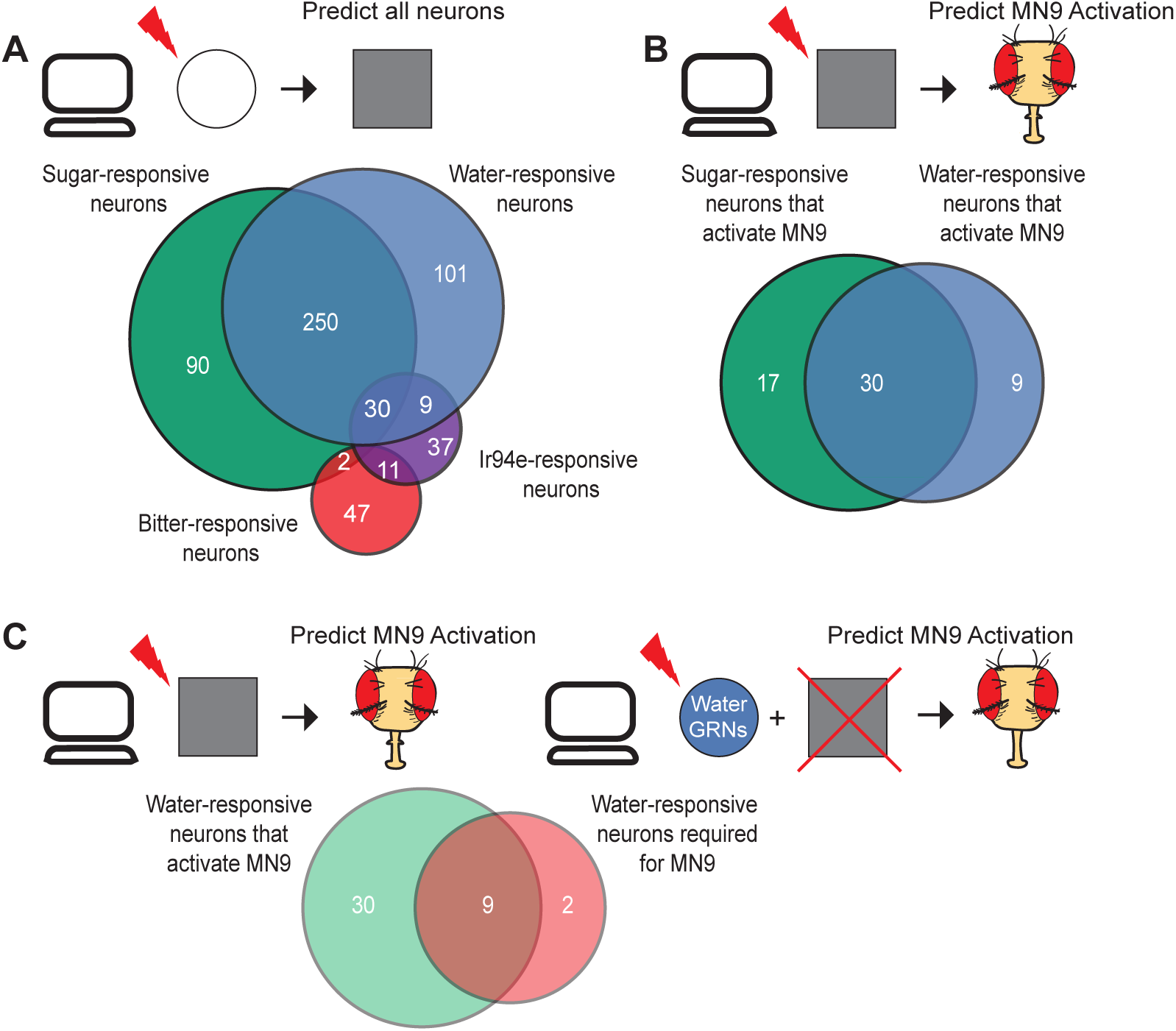
A comparision between predicted water, sugar, bitter and lr94e activation. A. Venn diagram showing the number and overlap of neurons that respond to sugar GRN (green) or water GRN (blue) activation that elicits 40 Hz MN9 firing, as well as bitter GRN (red) or lr94e GRN activation (purple) activated to reduce 40 Hz MN9 firing to 1 Hz. B. Venn diagram of the sugar-responsive neurons that are sufficient for MN9 activation and water-responsive neurons sufficient for MN9 activation. C. Venn diagram comparing predicted water-responsive neurons sufficient for MN9 and water-responsive neurons required (i.e., predicted to reduce MN9 firing >20%) for MN9 activity. GRNs are excluded from this analysis.

**Supplemental Figure 3.**
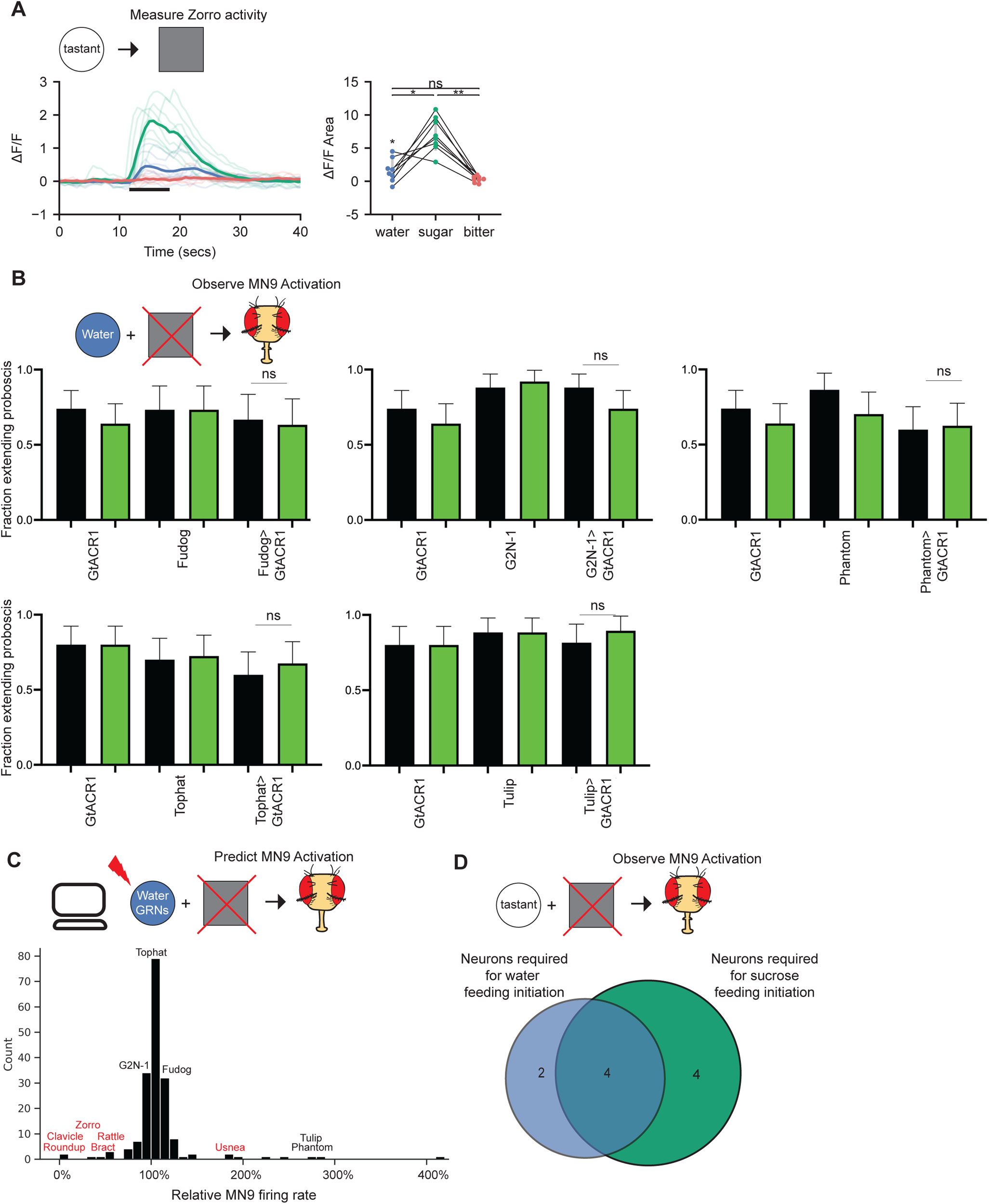
A comparision between predicted water silencing phenotypes and optogenetic silencing. A. Calcium responses of the second-order neuron Zorro to stimulation of the proboscis in pseudodessicated flies. B. The fraction of flies exhibiting proboscis extension response upon water stimulation. Green bars indicate green light condition. Mean +/- 95% confi­ dence intervals, Fisher’s exact test, n=40-50. ns: not significant. C. Histogram of water silencing predictions of the non-GRNs in Figure 4C at 160 Hz, with tested cell types labelled. Y-axis depicts the number of neurons in each bin. Cell types in red have an experimental water silencing phenotype. D. Comparison of neurons found to be required for feeding initiation to either water or 50 mM sucrose (Shiu, Sterne et al., 2022), when silenced with the anion channel rhodopsin GtACR1.

**Supplemental Figure 4.**
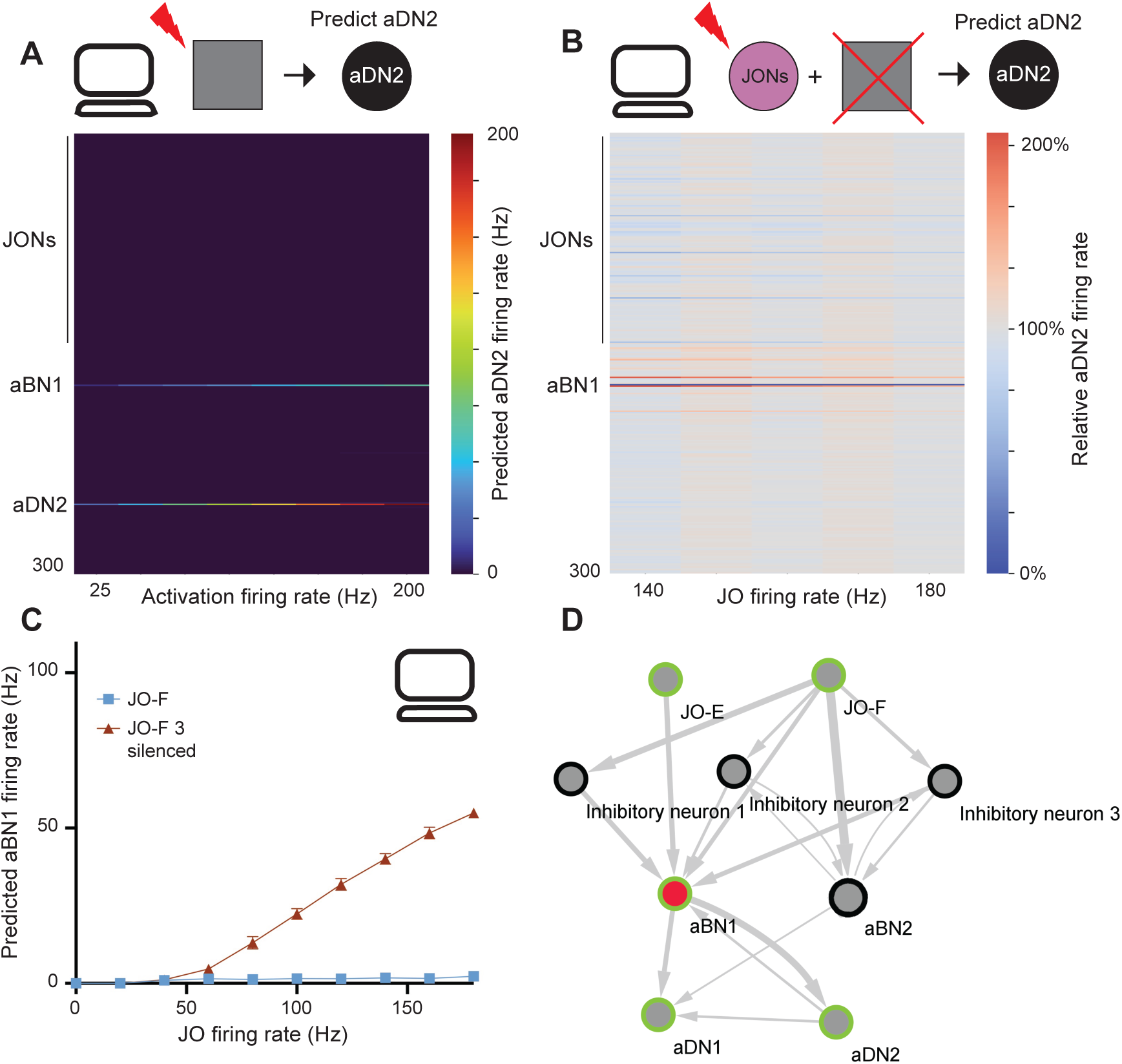
The computational model predicts neurons that elicit aDN2, and neurons that inhibit aBN1. A. Heatmap depicting the predicted aDN2 firing rate when the top 300 responsive neurons are activated at 25-200 Hz B. Heatmap depicting the change in aDN2 firing rate in response to activation of JOs at the specified firing rate, while individually silencing each of the top responsive neurons. C. Predicted aBN1 firing in response to JO-F activation at the specified rate. Red triangles, JO-F activation with co-silencing of three inhibitory neurons. D. Schematic of the circuit, with neurons sufficient for antenna! grooming in green, and required for antenna! grooming in red. Width of arrows indicates number of synaptic connections.

## Notes

### Competing Interest Statement

The authors have declared no competing interest.

## References

Baker, CA, McKellar C, Nern A, Dorkenwald S, Dickson BJ, Murthy M. Neural network organization for courtship-song feature detection in Drosophila. 2022. Current Biology. 32, 3317–3333.e7 doi: 10.1016/j.cub.2022.06.019

Bargmann CI. 2012. Beyond the connectome: How neuromodulators shape neural circuits. Bioessays 34:458–465. doi:10.1002/bies.201100185

Bargmann CI, Marder E. 2013. From the connectome to brain function. Nat Methods 10:483–490. doi:10.1038/nmeth.2451

Bates AS, Schlegel P, Roberts RJV, Drummond N, Tamimi IFM, Turnbull R, Zhao X, Marin EC, Popovici PD, Dhawan S, Jamasb A, Javier A, Serratosa Capdevila L, Li F, Rubin GM, Waddell S, Bock DD, Costa M, Jefferis GSXE. 2020. Complete Connectomic Reconstruction of Olfactory Projection Neurons in the Fly Brain. Current Biology 30:3183–3199.e6. doi:10.1016/j.cub.2020.06.042

Buhmann J, Sheridan A, Malin-Mayor C, Schlegel P, Gerhard S, Kazimiers T, Krause R, Nguyen TM, Heinrich L, Lee W-CA, Wilson R, Saalfeld S, Jefferis GSXE, Bock DD, Turaga SC, Cook M, Funke J. 2021. Automatic detection of synaptic partners in a whole-brain Drosophila electron microscopy data set. Nat Methods 18:771–774. doi:10.1038/s41592-021-01183-7

Cameron P, Hiroi M, Ngai J, Scott K. 2010. The molecular basis for water taste in Drosophila. Nature 465:91–95. doi:10.1038/nature09011

Chen T-W, Wardill TJ, Sun Y, Pulver SR, Renninger SL, Baohan A, Schreiter ER, Kerr RA, Orger MB, Jayaraman V, Looger LL, Svoboda K, Kim DS. 2013. Ultrasensitive fluorescent proteins for imaging neuronal activity. Nature 499:295–300. doi:10.1038/nature12354

Chen Z, Wang Q, Wang Z. 2010. The Amiloride-Sensitive Epithelial Na ^+^ Channel PPK28 Is Essential for *Drosophila* Gustatory Water Reception. J Neurosci 30:6247–6252. doi:10.1523/JNEUROSCI.0627-10.2010

Chu B, Chui V, Mann K, Gordon MD. 2014. Presynaptic Gain Control Drives Sweet and Bitter Taste Integration in Drosophila. Current Biology 24:1978–1984. doi:10.1016/j.cub.2014.07.020

Churgin MA, Lavrentovich D, Smith MA, Gao R, Boyden E, de Bivort B. 2021. Neural correlates of individual odor preference in *Drosophila* (preprint). Neuroscience. doi:10.1101/2021.12.24.474127

Dahanukar A, Foster K, van der Goes van Naters WM, Carlson JR. 2001. A Gr receptor is required for response to the sugar trehalose in taste neurons of Drosophila. Nat Neurosci 4:1182–1186. doi:10.1038/nn765

Dahanukar A, Lei Y-T, Kwon JY, Carlson JR. 2007. Two Gr Genes Underlie Sugar Reception in Drosophila. Neuron 56:503–516. doi:10.1016/j.neuron.2007.10.024

Dermietzel R, Spray DC. 1993. Gap junctions in the brain: where, what type, how many and why? Trends in Neurosciences 16:186–192. doi:10.1016/0166-2236(93)90151-B

Devineni, A, Sun B, Zhukovskaya A, Axel R. 2019. Acetic acid activates distinct taste pathways in Drosophila to elicit opposing, state-dependent feeding responses. eLife 8:e47677. doi: 10.7554/eLife.47677

Dionne H, Hibbard KL, Cavallaro A, Kao J-C, Rubin GM. 2018. Genetic Reagents for Making Split-GAL4 Lines in *Drosophila*. Genetics 209:31–35. doi:10.1534/genetics.118.300682

Dolan M-J, Frechter S, Bates AS, Dan C, Huoviala P, Roberts RJ, Schlegel P, Dhawan S, Tabano R, Dionne H, Christoforou C, Close K, Sutcliffe B, Giuliani B, Li F, Costa M, Ihrke G, Meissner GW, Bock DD, Aso Y, Rubin GM, Jefferis GS. 2019. Neurogenetic dissection of the Drosophila lateral horn reveals major outputs, diverse behavioural functions, and interactions with the mushroom body. eLife 8:e43079. doi:10.7554/eLife.43079

Dorkenwald S, McKellar CE, Macrina T, Kemnitz N, Lee K, Lu R, Wu J, Popovych S, Mitchell E, Nehoran B, Jia Z, Bae JA, Mu S, Ih D, Castro M, Ogedengbe O, Halageri A, Kuehner K, Sterling AR, Ashwood Z, Zung J, Brittain D, Collman F, Schneider-Mizell C, Jordan C, Silversmith W, Baker C, Deutsch D, Encarnacion-Rivera L, Kumar S, Burke A, Bland D, Gager J, Hebditch J, Koolman S, Moore M, Morejohn S, Silverman B, Willie K, Willie R, Yu S, Murthy M, Seung HS. 2022. FlyWire: online community for whole-brain connectomics. Nat Methods 19:119–128. doi:10.1038/s41592-021-01330-0

Eckstein N, Bates AS, Du M, Hartenstein V, Jefferis GSXE, Funke J. 2020. Neurotransmitter Classification from Electron Microscopy Images at Synaptic Sites in Drosophila (preprint). Neuroscience. doi:10.1101/2020.06.12.148775

Engert S, Sterne GR, Bock DD, Scott K. 2022. Drosophila gustatory projections are segregated by taste modality and connectivity. eLife 11:e78110. doi:10.7554/eLife.78110

Flavell SW, Gogolla N, Lovett-Barron M, Zelikowsky M. 2022. The emergence and influence of internal states. Neuron 110:2545–2570. doi:10.1016/j.neuron.2022.04.030

Flood TF, Iguchi S, Gorczyca M, White B, Ito K, Yoshihara M. 2013. A single pair of interneurons commands the Drosophila feeding motor program. Nature 499:83–87. doi:10.1038/nature12208

French AS, Sellier M-J, Ali Agha M, Guigue A, Chabaud M-A, Reeb PD, Mitra A, Grau Y, Soustelle L, Marion-Poll F. 2015. Dual Mechanism for Bitter Avoidance in *Drosophila*. J Neurosci 35:3990–4004. doi:10.1523/JNEUROSCI.1312-14.2015

Gerstner W, Kistler WM, Naud R, Paninski L. 2014. Neuronal dynamics: from single neurons to networks and models of cognition. Cambridge: Cambridge University Press.

Gordon MD, Scott K. 2009. Motor Control in a Drosophila Taste Circuit. Neuron 61:373–384. doi:10.1016/j.neuron.2008.12.033

Gouwens NW, Wilson RI. 2009. Signal Propagation in *Drosophila* Central Neurons. J Neurosci 29:6239–6249. doi:10.1523/JNEUROSCI.0764-09.2009

Hampel S, Eichler K, Yamada D, Bock DD, Kamikouchi A, Seeds AM. 2020. Distinct subpopulations of mechanosensory chordotonal organ neurons elicit grooming of the fruit fly antennae. eLife 9:e59976. doi:10.7554/eLife.59976

Hampel S, Franconville R, Simpson JH, Seeds AM. 2015. A neural command circuit for grooming movement control. eLife 4:e08758. doi:10.7554/eLife.08758

Hampel S, McKellar CE, Simpson JH, Seeds AM. 2017. Simultaneous activation of parallel sensory pathways promotes a grooming sequence in Drosophila. eLife 6:e28804. doi:10.7554/eLife.28804

Harris DT, Kallman BR, Mullaney BC, Scott K. 2015. Representations of Taste Modality in the Drosophila Brain. Neuron 86:1449–1460. doi:10.1016/j.neuron.2015.05.026

Hartenstein V, Omoto JJ, Ngo KT, Wong D, Kuert PA, Reichert H, Lovick JK, Younossi-Hartenstein A. 2018. Structure and development of the subesophageal zone of the *Drosophila* brain. I. Segmental architecture, compartmentalization, and lineage anatomy. J Comp Neurol 526:6–32. doi:10.1002/cne.24287

Heinrich L, Funke J, Pape C, Nunez-Iglesias J, Saalfeld S. 2018. Synaptic Cleft Segmentation in Non-isotropic Volume Electron Microscopy of the Complete Drosophila Brain In: Frangi AF, Schnabel JA, Davatzikos C, Alberola-López C, Fichtinger G, editors. Medical Image Computing and Computer Assisted Intervention – MICCAI 2018, Lecture Notes in Computer Science. Cham: Springer International Publishing. pp. 317–325. doi:10.1007/978-3-030-00934-2_36

Huang YC, Wang CT, Su TS, Kao KW, Lin YJ, Chuang CC, Chiang AS, Lo CC. 2018. A Single-Cell level and Connectome-Derived computational model of the Drosophila Brain Frontiers in Neuroinformatics 12:99. doi:10.3389/fninf.2018.00099

Inagaki HK, Ben-Tabou de-Leon S, Wong AM, Jagadish S, Ishimoto H, Barnea G, Kitamoto T, Axel R, Anderson DJ. 2012. Visualizing Neuromodulation In Vivo: TANGO-Mapping of Dopamine Signaling Reveals Appetite Control of Sugar Sensing. Cell 148:583–595. doi:10.1016/j.cell.2011.12.022

Inagaki HK, Panse KM, Anderson DJ. 2014. Independent, Reciprocal Neuromodulatory Control of Sweet and Bitter Taste Sensitivity during Starvation in Drosophila. Neuron 84:806–820. doi:10.1016/j.neuron.2014.09.032

Jaeger AH, Stanley M, Weiss ZF, Musso P-Y, Chan RC, Zhang H, Feldman-Kiss D, Gordon MD. 2018. A complex peripheral code for salt taste in Drosophila. eLife 7:e37167. doi:10.7554/eLife.37167

Kakaria KS, de Bivort BL. 2017. Ring Attractor Dynamics Emerge from a Spiking Model of the Entire Protocerebral Bridge. Front Behav Neurosci 11. doi:10.3389/fnbeh.2017.00008

Klapoetke NC, Murata Y, Kim SS, Pulver SR, Birdsey-Benson A, Cho YK, Morimoto TK, Chuong AS, Carpenter EJ, Tian Z, Wang J, Xie Y, Yan Z, Zhang Y, Chow BY, Surek B, Melkonian M, Jayaraman V, Constantine-Paton M, Wong GK-S, Boyden ES. 2014. Independent optical excitation of distinct neural populations. Nat Methods 11:338–346. doi:10.1038/nmeth.2836

Lappalainen JK, Tschopp, FD, Prakhya S, McGill M, Nern A, Shinomiya, K, Takemura, S, Gruntman E, Macke E, Turaga S. 2023. Connectome-constrained deep mechanistic networks predict neural responses across the fly visual system at single-neuron resolution (preprint). Neuroscience. doi:10.1101/2023.03.11.532232

Liu WW, Wilson RI. 2013. Glutamate is an inhibitory neurotransmitter in the *Drosophila* olfactory system. Proc Natl Acad Sci USA 110:10294–10299. doi:10.1073/pnas.1220560110

Mann K, Gordon MD, Scott K. 2013. A Pair of Interneurons Influences the Choice between Feeding and Locomotion in Drosophila. Neuron 79:754–765. doi:10.1016/j.neuron.2013.06.018

Marder E. 2012. Neuromodulation of Neuronal Circuits: Back to the Future. Neuron 76:1–11. doi:10.1016/j.neuron.2012.09.010

Marder E, Bucher D. 2007. Understanding Circuit Dynamics Using the Stomatogastric Nervous System of Lobsters and Crabs. Annu Rev Physiol 69:291–316. doi:10.1146/annurev.physiol.69.031905.161516

Marella S, Mann K, Scott K. 2012. Dopaminergic Modulation of Sucrose Acceptance Behavior in Drosophila. Neuron 73:941–950. doi:10.1016/j.neuron.2011.12.032

Marin EC, Büld L, Theiss M, Sarkissian T, Roberts RJV, Turnbull R, Tamimi IFM, Pleijzier MW, Laursen WJ, Drummond N, Schlegel P, Bates AS, Li F, Landgraf M, Costa M, Bock DD, Garrity PA, Jefferis GSXE. 2020. Connectomics Analysis Reveals First-, Second-, and Third-Order Thermosensory and Hygrosensory Neurons in the Adult Drosophila Brain. Current Biology 30:3167–3182.e4. doi:10.1016/j.cub.2020.06.028

McKellar CE, Siwanowicz I, Dickson BJ, Simpson JH. 2020. Controlling motor neurons of every muscle for fly proboscis reaching. eLife 9:e54978. doi:10.7554/eLife.54978

Meinertzhagen IA. 2018. Of what use is connectomics? A personal perspective on the *Drosophila* connectome. Journal of Experimental Biology 221:jeb164954. doi:10.1242/jeb.164954

Meunier N, Marion-Poll F, Rospars J-P, Tanimura T. 2003. Peripheral coding of bitter taste inDrosophila. J Neurobiol 56:139–152. doi:10.1002/neu.10235

Miyazaki T, Ito K. 2010. Neural architecture of the primary gustatory center of Drosophila melanogaster visualized with GAL4 and LexA enhancer-trap systems. J Comp Neurol 518:4147–4181. doi:10.1002/cne.22433

Miyazaki T, Lin T-Y, Ito K, Lee C-H, Stopfer M. 2015. A gustatory second-order neuron that connects sucrose-sensitive primary neurons and a distinct region of the gnathal ganglion in the *Drosophila* brain. Journal of Neurogenetics 29:144–155. doi:10.3109/01677063.2015.1054993

Mohammad F, Stewart JC, Ott S, Chlebikova K, Chua JY, Koh T-W, Ho J, Claridge-Chang A. 2017. Optogenetic inhibition of behavior with anion channelrhodopsins. Nat Methods 14:271–274. doi:10.1038/nmeth.4148

Montell C. 2021. *Drosophila* sensory receptors—a set of molecular Swiss Army Knives. Genetics 217:1–34. doi:10.1093/genetics/iyaa011

Münch D, Goldschmidt D, Ribeiro C. 2022. The neuronal logic of how internal states control food choice. Nature 607:747–755. doi:10.1038/s41586-022-04909-5

Nässel DR, Zandawala M. 2019. Recent advances in neuropeptide signaling in Drosophila, from genes to physiology and behavior. Progress in Neurobiology 179:101607. doi:10.1016/j.pneurobio.2019.02.003

Nusbaum MP, Blitz DM, Marder E. 2017. Functional consequences of neuropeptide and small-molecule co-transmission. Nat Rev Neurosci 18:389–403. doi:10.1038/nrn.2017.56

Ohyama T, Schneider-Mizell CM, Fetter RD, Aleman JV, Franconville R, Rivera-Alba M, Mensh BD, Branson KM, Simpson JH, Truman JW, Cardona A, Zlatic M. 2015. A multilevel multimodal circuit enhances action selection in Drosophila. Nature 520:633–639. doi:10.1038/nature14297

Paul MM, Pauli M, Ehmann N, Hallermann S, Sauer M, Kittel RJ, Heckmann M. 2015. Bruchpilot and Synaptotagmin collaborate to drive rapid glutamate release and active zone differentiation Frontiers in Cellular Neuroscience 9:29. doi: 10.3389/fncel.2015.00029

Pisokas I, Heinze S, Webb B. 2020. The head direction circuit of two insect species. eLife 9:e53985. doi:10.7554/eLife.53985

Rozental R, Giaume C, Spray DC. 2000. Gap junctions in the nervous system. Brain Research Reviews 32:11–15. doi:10.1016/S0165-0173(99)00095-8

Scheffer LK, Meinertzhagen IA. 2021. A connectome is not enough – what is still needed to understand the brain of *Drosophila* ? Journal of Experimental Biology 224:jeb242740. doi:10.1242/jeb.242740

Scheffer LK, Xu CS, Januszewski M, Lu Z, Takemura Shin-ya, Hayworth KJ, Huang GB, Shinomiya K, Maitlin-Shepard J, Berg S, Clements J, Hubbard PM, Katz WT, Umayam L, Zhao T, Ackerman D, Blakely T, Bogovic J, Dolafi T, Kainmueller D, Kawase T, Khairy KA, Leavitt L, Li PH, Lindsey L, Neubarth N, Olbris DJ, Otsuna H, Trautman ET, Ito M, Bates AS, Goldammer J, Wolff T, Svirskas R, Schlegel P, Neace E, Knecht CJ, Alvarado CX, Bailey DA, Ballinger S, Borycz JA, Canino BS, Cheatham N, Cook M, Dreher M, Duclos O, Eubanks B, Fairbanks K, Finley S, Forknall N, Francis A, Hopkins GP, Joyce EM, Kim S, Kirk NA, Kovalyak J, Lauchie SA, Lohff A, Maldonado C, Manley EA, McLin S, Mooney C, Ndama M, Ogundeyi O, Okeoma N, Ordish C, Padilla N, Patrick CM, Paterson T, Phillips EE, Phillips EM, Rampally N, Ribeiro C, Robertson MK, Rymer JT, Ryan SM, Sammons M, Scott AK, Scott AL, Shinomiya A, Smith C, Smith K, Smith NL, Sobeski MA, Suleiman A, Swift J, Takemura Satoko, Talebi I, Tarnogorska D, Tenshaw E, Tokhi T, Walsh JJ, Yang T, Horne JA, Li F, Parekh R, Rivlin PK, Jayaraman V, Costa M, Jefferis GS, Ito K, Saalfeld S, George R, Meinertzhagen IA, Rubin GM, Hess HF, Jain V, Plaza SM. 2020. A connectome and analysis of the adult Drosophila central brain. eLife 9:e57443. doi:10.7554/eLife.57443

Schlegel P, Bates AS, Stürner T, Jagannathan SR, Drummond N, Hsu J, Capdevila LS, Javier A, Marin EC, Barth-Maron A, Tamimi IFM, Li F, Rubin GM, Plaza SM, Costa M, Jefferis GSXE. 2020. Information flow, cell types and stereotypy in a full olfactory connectome (preprint). Neuroscience. doi:10.1101/2020.12.15.401257

Schwarz O, Bohra AA, Liu X, Reichert H, VijayRaghavan K, Pielage J. 2017. Motor control of Drosophila feeding behavior. eLife 6:e19892. doi:10.7554/eLife.19892

Scott K, Brady R, Cravchik A, Morozov P, Rzhetsky A, Zuker C, Axel R. 2001. A Chemosensory Gene Family Encoding Candidate Gustatory and Olfactory Receptors in Drosophila. Cell 104:661–673. doi:10.1016/S0092-8674(01)00263-X

Seeds AM, Ravbar P, Chung P, Hampel S, Midgley FM, Mensh BD, Simpson JH. 2014. A suppression hierarchy among competing motor programs drives sequential grooming in Drosophila. eLife 3:e02951. doi:10.7554/eLife.02951

Shiu PK, Sterne GR, Engert S, Dickson BJ, Scott K. 2022. Taste quality and hunger interactions in a feeding sensorimotor circuit. eLife 11:e79887. doi:10.7554/eLife.79887

Sterne GR, Otsuna H, Dickson BJ, Scott K. 2021. Classification and genetic targeting of cell types in the primary taste and premotor center of the adult Drosophila brain. eLife 10:e71679. doi:10.7554/eLife.71679

Stocker RF. 1994. The organization of the chemosensory system in Drosophila melanogaster: a review. Cell Tissue Res 275:3–26. doi:10.1007/BF00305372

Stocker RF, Schorderet M. 1981. Cobalt filling of sensory projections from internal and external mouthparts in Drosophila. Cell Tissue Res 216. doi:10.1007/BF00238648

Thorne N, Chromey C, Bray S, Amrein H. 2004. Taste Perception and Coding in Drosophila. Current Biology 14:1065–1079. doi:10.1016/j.cub.2004.05.019

Wang Z, Singhvi A, Kong P, Scott K. 2004. Taste Representations in the Drosophila Brain. Cell 117:981–991. doi:10.1016/j.cell.2004.06.011

Winding M, Pedigo BD, Barnes CL, Patsolic HG, Park Y, Kazimiers T, Fushiki A, Andrade IV, Khandelwal A, Valdes-Aleman J, Li F, Randel N, Barsotti E, Correia A, Fetter RD, Hartenstein V, Priebe CE, Vogelstein JT, Cardona A, Zlatic M. 2023. The connectome of an insect brain Science 379:eadd9330. doi: 10.1126/science.add9330

Yetman S, Pollack GS. 1987. Proboscis extension in the blowfly: directional responses to stimulation of identified chemosensitive hairs. J Comp Physiol 160:367–374. doi:10.1007/BF00613026

Zhang N, Guo L, Simpson JH. 2020. Spatial Comparisons of Mechanosensory Information Govern the Grooming Sequence in Drosophila. Current Biology 30:988–1001.e4. doi:10.1016/j.cub.2020.01.045

Zheng Z, Lauritzen JS, Perlman E, Robinson CG, Nichols M, Milkie D, Torrens O, Price J, Fisher CB, Sharifi N, Calle-Schuler SA, Kmecova L, Ali IJ, Karsh B, Trautman ET, Bogovic JA, Hanslovsky P, Jefferis GSXE, Kazhdan M, Khairy K, Saalfeld S, Fetter RD, Bock DD. 2018. A Complete Electron Microscopy Volume of the Brain of Adult Drosophila melanogaster. Cell 174:730–743.e22. doi:10.1016/j.cell.2018.06.019

